# Genome-informed microscopy reveals infections of uncultivated carbon-fixing archaea by lytic viruses in Earth’s crust

**DOI:** 10.1101/2020.07.22.215848

**Authors:** Janina Rahlff, Victoria Turzynski, Sarah P. Esser, Indra Monsees, Till L.V. Bornemann, Perla Abigail Figueroa-Gonzalez, Frederik Schulz, Tanja Woyke, Andreas Klingl, Cristina Moraru, Alexander J. Probst

**Affiliations:** University of Duisburg-Essen, Department of Chemistry, Environmental Microbiology and Biotechnology (EMB), Group for Aquatic Microbial Ecology, Universitätsstraße 5, 45141 Essen, Germany; DOE Joint Genome Institute, 1 Cyclotron Rd, Berkeley, CA, 94720, USA; Plant Development & Electron Microscopy, Biocenter LMU Munich, Großhaderner Str. 2-4, 82152 Planegg-Martinsried, Germany; Institute for Chemistry and Biology of the Marine Environment (ICBM), Carl-von-Ossietzky-University Oldenburg, PO Box 2503, Carl-von-Ossietzky-Straße 9-11, 26111, Oldenburg, Germany

## Abstract

The continental subsurface houses a major portion of life’s abundance and diversity, yet little is known about viruses infecting microbes that reside there. Here, we used a combination of metagenomics and genome-informed microscopy to show that highly abundant carbon-fixing organisms of the uncultivated genus Candidatus Altiarchaeum are frequent targets of previously unrecognized viruses in the deep subsurface. Analysis of CRISPR spacer matches displayed resistances of Ca. Altiarchaea against eight predicted viral clades, which showed genomic relatedness across continents but little similarity to previously identified viruses. Based on metagenomic information, we tagged and imaged a putatively viral genome rich in protospacers using fluorescence microscopy. Virus-targeted genomeFISH revealed a lytic lifestyle of the respective virus and challenges previous predictions that lysogeny prevails as the dominant viral lifestyle in the subsurface. CRISPR development over time and imaging of 18 samples from one subsurface ecosystem suggest a sophisticated interplay of viral diversification and adapting CRISPR-mediated resistances of Ca. Altiarchaeum. We conclude that infections of primary producers with lytic viruses followed by cell lysis potentially jump-start heterotrophic carbon cycling in these subsurface ecosystems.

## INTRODUCTION

Earth’s continental subsurface harbours 2-6 × 10^29^ prokaryotic cells [1, 2], which represent a major component of life’s diversity on our planet [3, 4]. Among these organisms are some of the most enigmatic archaea, including Aigarchaeota, Asgard archaea, Altiarchaeota and members of the DPANN radiation [5–8]. Although the ecology and diversity of subsurface microorganisms has been under investigation in several studies, the fundamental question relating to how microbial diversity and composition in the deep subsurface change with virus infection remains mostly unanswered. Viruses have long been recognized as major drivers of microbial diversification [9], yet little is known about their lifestyle, activity and impact on oligotrophic subsurface ecosystems. Recent evidence demonstrated high numbers of virus-cell ratios in marine subsurface sediments [10], suggesting ongoing viral proliferation in the deep biosphere. In addition, pronounced morphological diversity of bacteriophages with presumably lytic representatives has been found in granitic groundwater of up to 450 m depth [11], and might be the result of recombination events, horizontal gene transfer and lysogeny known to shape microbial communities of the subsurface [12]. The recent recovery of two novel bacteriophage genera with lytic genes from groundwater highlights the potential of subsurface environments for being huge reservoirs of previously unknown viruses [13]. Furthermore, a study on predominant *Halanaerobium* spp. from anthropogenic subsurface communities (hydraulically fractured wells) suggested long-term host-virus dynamics, extensive viral predation and adaptive host immunity based on clustered regularly interspaced short palindromic repeats (CRISPR) spacer to protospacer matches [14]. CRISPR systems function as defense mechanisms for bacteria and archaea against mobile genetic elements (MGEs), including viruses [15]. The CRISPR locus is usually flanked by *cas* genes and interspaced by short variable DNA sequences termed spacers [15] previously acquired from invading MGEs. The diversification of CRISPR-Cas immunity in the host over geographical distances and time due to preceding viral infections and protospacer mutations has been well-documented, e.g., for *Sulfolobus islandicus* [16].

Oligotrophic anaerobic subsurface environments can be populated by a variety of different microorganisms, some of them belonging to the phylum Altiarchaeota [7, 17]. In fact, these organisms can reach high abundances in their ecosystems with up to 70% of the total community in the aquifer or with up to 95% within the biofilm (BF) they form [18, 19]. Members of the genus *Ca.* Altiarchaeum–*Ca.* A. hamiconexum being the best-studied representative [7]–occur in anoxic subsurface environments around the globe [20, 21] and fix carbon via the reductive acetyl-CoA (Wood-Ljungdahl) pathway [7]. In certain ecosystems, *Ca.* Altiarchaea form nearly pure BFs, which are kept together by filamentous cell surface appendages called “*hami*“ (singular: *hamus*) [22]. Studies to date showed symbiotic relationships of *Ca.* Altiarchaea with bacterial partners *Thiothrix* sp. [23], and *Sulfuricurvum* sp. [24], but also a co-occurrence with the episymbiont *Ca.* Huberiarchaeum crystalense, belonging to the DPANN clade, has been recently reported [17, 25]. A single transmission electron micrograph and the presence of CRISPR systems led to speculations on the existence of *Ca.* Altiarchaeum viruses in the subsurface [26]. However, mesophilic archaeal viruses from the deep terrestrial subsurface remain highly enigmatic, despite the fact that mesophilic archaeal genomes contain more MGEs than their thermophilic counterparts [27]. The knowledge gap on archaeal viruses is fostered by a lack of their genome entries in public databases [28], missing marker genes for viruses [29] and a bias towards viruses related to economical, medical or biotechnological activities [30]. In addition, only ~150 archaeal viruses have been isolated and described to date [31]. Recent exhaustive metagenomic surveys aided the discovery of novel archaeal viruses [32] from multiple ecosystems, including the ocean [33, 34], hot springs [35–37] and soils [38, 39], and eventually allowed targeting and visualization of an uncultivated virus based on its genome [40].

Due to their world-wide distribution and high abundance as the main primary producer in certain continental subsurface ecosystems, *Ca.* Altiarchaea represent the ideal model genus for studying viruses and their infection mechanisms of mesophilic microorganisms in the subsurface. This is especially relevant because the extent to which lytic infections occur in the continental subsurface is unknown, and lysogeny is assumed to be the predominant viral strategy [41]. Consequently, we used metagenomics to predict viruses that infect *Ca.* Altiarchaea in subsurface ecosystems at four different sites across three continents. The most abundant putative virus, which showed little homologies with sequences in public databases and thus carried no viral hallmark genes, was visualized and characterized using genome-informed microscopy, providing novel insights into its viral lifestyle. Our analyses further demonstrate the diversification of CRISPR systems of *Ca.* Altiarchaea along with a decline in virus abundances over six years.

## MATERIAL AND METHODS

### Mining public metagenomes for Altiarchaeota

To get an overview of the global distribution of Altiarchaeota, we searched metagenomes in the IMG/M database [42] (database accessed in July 2018) for Altiarchaeota contigs using DIAMOND blastp (v0.9.22) [43] with the putative *hamus* subunit (NCBI accession no CEG12198.1) as a query and an e-value and length cut-off of 1e-10 and 300 amino acids, respectively. The Altiarchaeota distribution based on 16S rRNA gene sequences was obtained from the SILVA SSU Parc database [44] based on all 16S rRNA genes classified as Altiarchaeota and for which geographic information was available (July 2018).

To investigate Altiarchaeota-virus relationships, we explored ecosystems where Altiarchaeota comprised the majority of the community. Therefore, metagenomic data from a microbial mat growing in the sulfidic groundwater-fed Alpena County Library Fountain (ACLF) (Alpena, MI, USA) were obtained from NCBI Sequence Read Archive (SRA) repository [45] as were metagenome samples from Horonobe Underground Research Laboratory at 140 and 250 m depth (HURL, Hokkaido, Japan) [21]. The analysis was complemented with three metagenomic datasets from CO_2_-enriched groundwater erupted from a cold-water Geyser (Andernach, Middle Rhine Valley, Germany) [46], and the metagenome of a sulfidic spring Mühlbacher Schwefelquelle Isling (MSI) in Regensburg, Germany [7]. All BioProject, BioSample accessions and information on sampling of MSI for various experiments are provided in Table S1 and S2, respectively.

### Re-sampling of MSI for virus-targeted genomeFISH and metagenomic sequencing

BF samples were collected as previously described [19] from the 36.5 m deep, cold (~10 °C), sulfidic spring MSI in Regensburg, Germany (N 48° 59.142, E 012° 07.636) in January 2019, for which further geological description has been reported elsewhere [47]. Environmental parameters of the sulfidic spring were extensively investigated previously [18, 23] and remained almost constant over years. For virus-targeted genomeFISH, BF samples were fixed by addition of formaldehyde (3% v/v) and incubation at room temperature for one hour. For investigating infection stages, BF flocks were gently separated from several bigger flocks by using a pipette tip finally yielding 18 smaller flocks (range 170 – 5 870 μm^2^). BF flocks for all microscopy experiments were subsequently washed three times in 1x phosphate buffered saline (PBS, pH 7.4) followed by dehydration via an ethanol gradient (50%, 70% v/v, and absolute ethanol, 10 min each). Samples were stored in absolute ethanol at −20 °C until further processing.

Three types of samples were collected for metagenomics: i) BF flocks; ii) the planktonic community (>0.1 μm pore-size fraction); and iii) the viruses and lysed cells (<0.1 μm pore-size fraction). Sampling of Altiarchaeota BF flocks for DNA extraction and metagenomic sequencing from the sulfidic spring (MSI) was performed in October 2018. For sampling the unfiltered planktonic microbial community, 70 L of groundwater were filtered onto a 0.1 μm pore-size PTFE membrane filter (Merck Millipore, Darmstadt, Germany). The flow-through was collected in a sterilized container, and a final concentration of 1 mg L^−1^ of iron (III) chloride (Carl Roth, Karlsruhe, Germany) was applied for chemical flocculation for 30 minutes [48]. Flocculates were filtered onto 5 × 0.2 μm membrane filters (<0.1 μm fraction). DNA was extracted directly from collected BF samples using the RNeasy^®^ PowerBiofilm Kit (Qiagen, Hilden, Germany) using a DNA-conform workflow. DNA from the 0.1 μm and pooled 0.2 μm membrane filters with iron flocculates was extracted using PowerMax Soil DNA Extraction Kit (Qiagen, Hilden, Germany), DNA was precipitated overnight and cleaned with 70% ethanol. Shotgun metagenome sequencing was conducted within the Census of Deep Life Sequencing call 2018 and performed using the Illumina HiSeq platform at the Marine Biological Laboratory, Woods Hole, MA, USA.

### Detection of viral genomes in metagenomes

Raw shotgun sequencing reads were trimmed and quality-filtered using bbduk (https://github.com/BioInfoTools/BBMap/blob/master/sh/bbduk.sh) and Sickle [49]. Read assembly was conducted by using metaSPADes v.3.10 [50] unless stated otherwise. Scaffolds <1 kb length were excluded from further analysis. Genes were predicted using prodigal [51] (meta mode) and functional annotations were determined by using DIAMOND [43] against UniRef100 [52]. Altiarchaeota genomes were retrieved from public datasets as described previously [46] and further cleaned or re-binned using GC, coverage and taxonomy information [53], which was necessary for all genomes except for Altiarchaeota from MSI_BF_2012 (Table S1). Viral scaffolds >3 kb were identified by applying a combination of tools as presented in Figure S1A. Predicted viruses were classified into viruses and putative viruses according to the classification system presented in Figure S1B. Viral scaffolds were subsequently checked for mini-CRISPR arrays using default settings of CRISPRCasFinder [54] as some archaeal viruses can bear mini-CRISPR arrays with 1-2 spacers having likely a role in interviral conflicts [55, 56].

### CRISPR-Cas analysis of Altiarchaeota genomes

*Cas* genes and direct repeat (DR) sequences were identified in binned Altiarchaeota genomes via CRISPRCasFinder [54] and genes were additionally confirmed via searches against UniRef100 [52]. CRISPR DR sequences were tested for formation of secondary structures using RNAfold [57] and checked against the CRISPRmap database (v2.1.3-2014) [58] for formation of known motifs. The consensus DR sequence was used in both possible orientations to extract host-specific spacers from raw reads by using MetaCRAST [59] with Cd-hit [60] clustering at 99% identity. Spacers were filtered for minimum and maximum lengths of 20 and 60 nucleotides, respectively. Only spacers that were present on a read that contained at least one complete DR sequence with an exact match to the template were considered. Finally, spacers were further clustered with Cd-hit [60] at 99% identity and matched to viral protospacers on the compiled output of viral identification tools using the blastn --short algorithm with a 80% similarity threshold. For comparing spacer dynamics (total abundance, diversity and matches to Altivir_1_MSI and Altivir_2_MSI genomes) of 2012 and 2018 samples from MSI, all spacers of the four respective MSI samples were clustered with Cd-hit [60] at 99% identity and representative sequences of each cluster were matched to representative Altivir_1_MSI and Altivir_2_MSI genomes from the BF of 2012 (Table 1). All data arising from spacer counts were normalized by genome abundance of the respective Altiarchaeota genome.

**Table 1:**
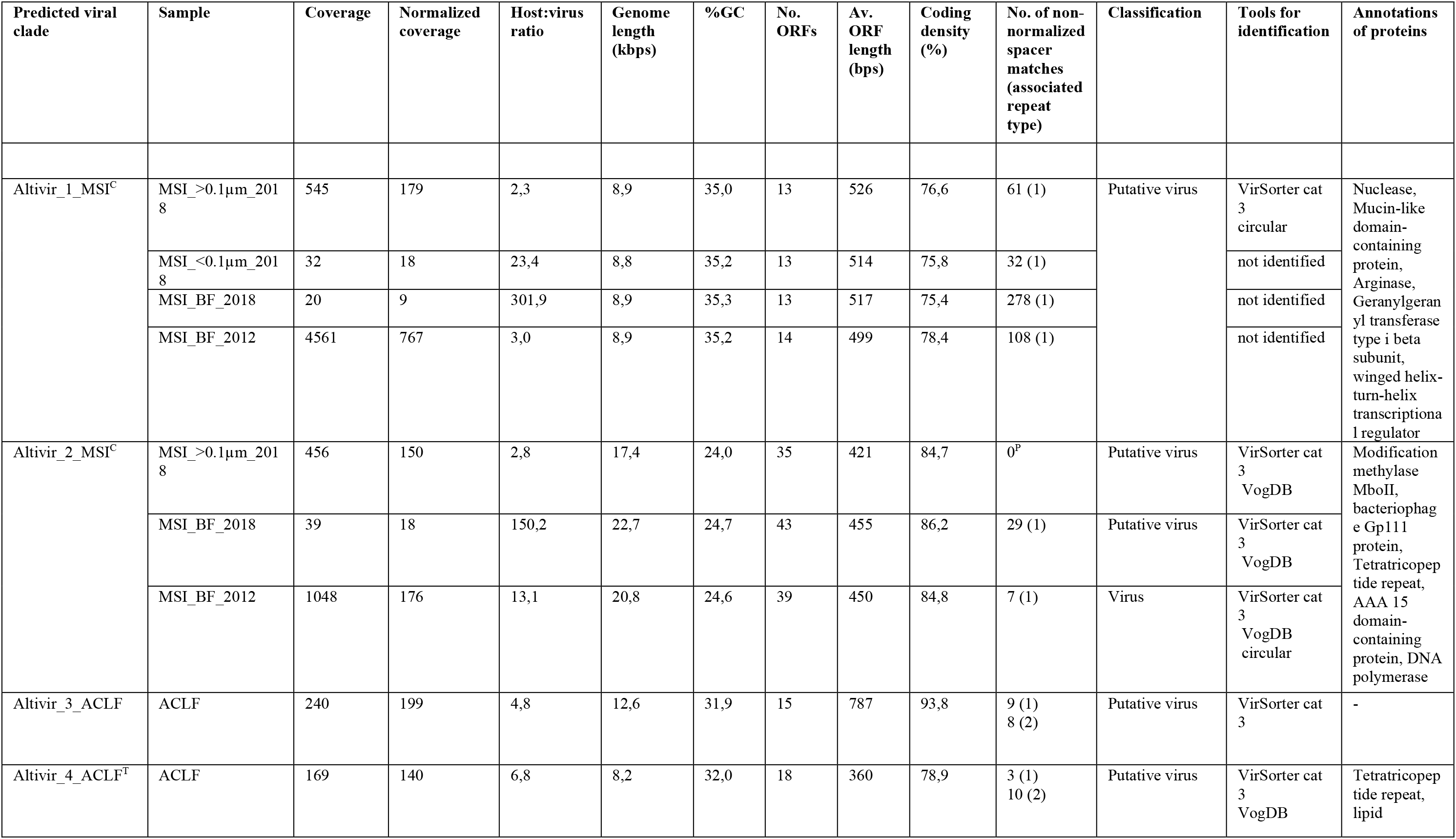
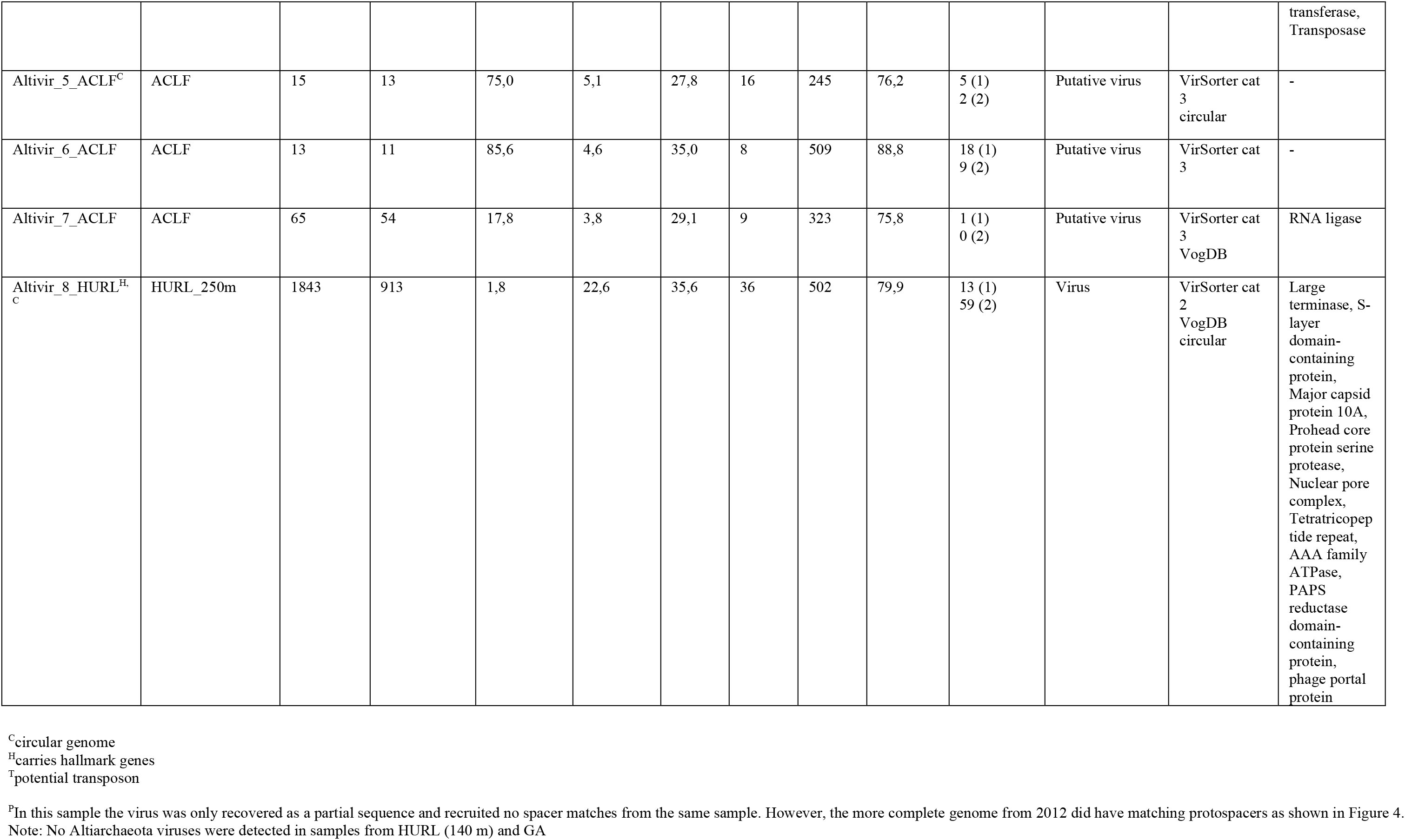
Characteristics of potential viruses of Altiarchaeota detected in subsurface ecosystems. Viruses were clustered at genus level and based on intergenomic similarity (Figure 2, Figure S4 and S5).

### Annotation and clustering of viral genomes

Coding sequences of predicted viral scaffolds were identified using prodigal (meta mode) [51] and respective proteins were annotated via DIAMOND search [43] against UniRef100 [52] by applying HHpred [61, 62] with an e-value cut-off of 10^−6^ against the following databases: PDB_mmCIFC70_4_Feb, Pfam-A v.32.0, NCBI_Conserved_Domains_v3.16 and TIGRFAMs v15.0 database. For the two most abundant predicted viruses Altivir_1_MSI and Altivir_8_HURL we additionally applied DELTA-BLAST [63] searches against NCBI’s non-redundant protein sequences (nr) and used PHMMR [64] against reference proteomes. Predicted viral scaffolds carrying multiple hits against existing bacterial genomes in UniRef100 [52] and no viral hallmark genes or carrying extensive CRISPR arrays (detected as false positives) were removed from further analyses.

Clustering of entire viral genomes on nucleic-acid level was performed with VICTOR [65] using the Genome-BLAST Distance Phylogeny method [66] with distance formula d0 and OPTSIL clustering [67]. A separate virus clustering was performed using vConTACT2 [68, 69] applying the database ‘ProkaryoticViralRefSeq94’ [70] and visualization of the viral network was performed in Cytoscape version 3.7.2 [71]. Intergenomic similarities between viral genomes and the corresponding heatmap were calculated using VIRIDIC accessible through http://viridic.icbm.de with default settings [72]. In order to further investigate the genetic relationship between the Altiarchaeota viruses, proteins of all viral genomes were clustered by first performing an all against all blastp with an e-value cut-off of 10^−5^ and a bitscore threshold of 50. Then, the results were loaded into the mcl program with the parameters “ -l --abc”. The genomic maps were plotted using the genoPlotR v.0.8.9 [73] package of the R programming environment [74].

### Relative abundance of viruses and hosts

Abundance of hosts and viruses was determined via read mapping respective genomes using Bowtie2 [75] in sensitive mode followed by mismatch filtering for host genomes (2%, depending on read length). Abundances were normalized to the total number of base pairs (bp) sequenced of each sample scaling to the sample with lowest counts, which is herein referred to as relative abundance or normalized coverage. Rank abundance curves were built based on the abundance of scaffolds carrying ribosomal protein S3 (*rpS3*) sequences predicted in metagenomic assemblies after running prodigal (meta mode) [51] and annotation [43] against UniRef100 [52]. For normalized rank abundance curves based on sequencing depth for the ecosystem MSI, we only considered *rpS3* sequences with a coverage >2.4 after normalization as this value represents the lowest coverage of any assembled *rpS3* sequence in the MSI dataset.

### Design, synthesis and chemical labeling of gene probes

To target Altivir_1_MSI, a 8.9 kb putative viral genome recovered from the MSI_BF_2012 metagenomic dataset, eleven dsDNA polynucleotide probes of 300 nucleotides length were designed using genePROBER (gene-prober.icbm.de/), according to [76]. In addition, a single 300 bp fragment of a *Metallosphaera* turreted icosahedral virus strain MTIV1 (NCBI accession no. MF443783.1) was designed as a negative control probe, because the *Metallosphaera* sp. virus was not detected in the metagenome of MSI_BF_2012. Sequences for all probes are given in Table S3.

The probes were synthesized as described in [76]. Shortly, all polynucleotides were chemically synthesized as gBlocks^®^ Gene Fragments (Integrated DNA Technologies, San Jose, CA, USA), reconstituted in 5 mM Tris-HCl pH 8.0, 1 mM EDTA pH 8.0 and then labeled with the ULYSIS™ Alexa Fluor™ 594 nucleic acid labeling kit (Thermo Fisher Scientific, MA, USA). Two μg of either an equimolar mixture of the eleven Altivir_1_MSI polynucleotides or of the single negative control polynucleotide were used in a single labeling reaction and then purified using NucAway Spin Columns (Thermo Fisher Scientific, USA). Before being used as probes for virus-targeted genome fluorescence *in situ* hybridization (genomeFISH), the labeled probes were measured spectrophotometrically using a NanoDrop™ (Thermo Fisher Scientific, USA). The calculated labeling efficiency was 9.16 and 16.6 dyes per base for Altivir_1_MSI probe and negative control probe, respectively.

### Virus-targeted genomeFISH of MSI biofilms

Virus-targeted genomeFISH was performed according to the direct-geneFISH protocol [77], with the modifications detailed further. In brief, Altiarchaeota BF, dehydrated in an ethanol series, were carefully placed in the middle of a press-to-seal silicone isolator (Sigma-Aldrich Chemie GmbH, Taufkirchen, Germany) mounted on Superfrost^®^ Plus slides (Electron Microscopy Sciences, Hatfield, USA), and subsequently air-dried. Because Altiarchaeota lack the typical archaeal S-layer as outer sheath [78] and hence are more prone to membrane disintegration, no extra permeabilization step was required. Different formamide concentrations (20%, 30%, 50%) were tested to exclude false positive hybridization signals. In the final assay, 20% of formamide was used in the hybridization buffer that contained 5x SSC buffer (saline sodium citrate, pH 7.0), 20% (w/v) dextran sulfate, 20 mM EDTA, 0.25 mg mL^− 1^ sheared salmon sperm DNA, 0.25 mg mL^−1^ yeast RNA, 1x blocking reagent, 0.1% (v/v) sodium dodecyl sulfate and nuclease-free water. The final NaCl concentration of 0.225M in the washing buffer corresponded to the 20% formamide in the hybridization buffer. As rRNA probes, the dual-Atto488-labeled probes NON338 [79] and a SM1-Euryarchaeon-specific probe “SMARCH714” [80] were used. A volume of 45 μl hybridization mixture was used, with a final gene probe concentration of 330 pg μL^−1^ (30 pg μL^−1^ for each polynucleotide) and rRNA probe final concentration of 1 pmol μL^−1^. In the hybridization chamber, 30 ml of formamide-water solution were added to keep a humid atmosphere. The denaturation and hybridization times were 30 min and 3 hours, respectively. The post hybridization washing buffer contained 20 mM Tris-HCl (pH 8.0), 5 mM EDTA (pH 8.0), nuclease-free water, 0.01 % SDS, and 0.225 M NaCl. A second washing step with 1x PBS (pH 7.4) for 20 min was performed. Then, the slides were transferred for one minute into molecular grade water and quickly rinsed in absolute ethanol. For staining, we used 15 μL of a mixture of 4’,6-diamidin-2-phenylindole (4 μg mL^−1^) in SlowFade Gold Antifade Mounting medium (both Thermo Fisher Scientific, Waltham, MA, USA). All solutions and buffers for virus-targeted genomeFISH experiments were prepared with molecular grade water (Carl Roth, Karlsruhe, Germany). For the experiment with individual flocks, 17 out of 18 flocks were treated with the Altivir_1_MSI probe and one flock with the negative control probe.

The BF material was examined and imaged with an Axio Imager M2m epifluorescence microscope equipped with an Axio Cam MRm and a Zen 2 Pro software (Carl Zeiss Microscopy GmbH, Jena, Germany). Channel mode visualization was performed by using the 110x/1.3 oil objective EC-Plan NEOFLUAR (Carl Zeiss Microscopy GmbH) and three different filter sets from Carl Zeiss: 49 DAPI for visualizing Altiarchaeota cells, 64 HE mPlum for the detection of viral infections, and 09 for achieving 16S rRNA signals. Enumeration of cells and viral signals was performed manually. Viral signals were categorized into three major groups, i.e., viral adsorption on host cells, advanced infections and viral bursts.

### Structured illumination microscopy

For structured illumination microscopy, virus-targeted genomeFISH was carried out on a BF flock mounted on a cover slip (thickness No. 1.5H, Paul Marienfeld GmbH & Co. KG, Lauda-Königshofen, Germany) and processed as stated above except that no DAPI was added to the mounting medium. Samples were analyzed using an inverted epifluorescence microscope (Zeiss ELYRA PS.1) equipped with an α-Plan-Apochromat 100x/1.46 oil DIC M27 Elyra objective, F Set 77 He filter, and a Zen black edition software, all obtained from the manufacturer Carl Zeiss Microscopy GmbH.

### Transmission electron microscopy

BF flocks from MSI were pre-fixed in glutaraldehyde (Carl Roth, Karlsruhe, Germany) to a final concentration of 2.5% (v/v) and physically fixed via high-pressure freezing followed by freeze substitution, which was carried out with 0.2% osmium tetroxide, 0.25% uranyl acetate and 9.3% water in acetone as described previously [81]. After embedding in Epon resin and polymerization for 72 h, the samples were ultrathin sectioned and post-stained with 1% lead citrate for two minutes. Transmission electron microscopy was carried out on a Zeiss EM 912 (Zeiss, Oberkochen, Germany) with an integrated OMEGA-filter at 80 kV in the zero-loss mode. Imaging was done using a 2k × 2k pixel slow-scan CCD camera (TRS Tröndle Restlichtverstärkersysteme, Moorenweis, Germany).

## RESULTS

### Globally distributed *Ca.* Altiarchaea have complex CRISPR systems with conserved DR sequences

Screening of 16S ribosomal RNA (rRNA) datasets and metagenomes within IMG [42] confirmed a global distribution of organisms belonging to the phylum Altiarchaeota (Figure S1). We performed metagenomic analyses of four terrestrial subsurface ecosystems that showed high abundance of the genus *Ca. Altiarchaeum* (Table S1), previously also termed Alti-1 [20], ranging from 37 to 352 m below ground. These ecosystems included i) an anoxic aquifer accessible through an artesian well (Mühlbacher Schwefelquelle, Isling, MSI) [24] sampled in 2012 and 2018 and ii) a high-CO_2_ geyser (Geyser Andernach, GA) [46] both located in Germany, iii) a sulfidic spring in the US (Alpena County Library Fountain, ACLF) [45], and iv) a deep underground laboratory in Japan (Horonobe Underground Research Laboratory, HURL) at 140 and 250 m depth [21]. All eight genomes of *Ca.* Altiarchaeum (Table S1) carried genetic information for Type I-B-CRISPR-Cas immunity including proteins Cas5, Cas7, Cas8a. Proteins of a Type III CRISPR-Cas immunity were found at the ACLF, HURL, and MSI site, including Repeat Associated Mysterious Proteins (RAMP, Cmr) of Type III-B and III-C, and Csm proteins of the Type III-A system (Table S4). Confidence in binning CRISPR arrays and assigning DR sequences to *Ca.* Altiarchaea arose from the 16-146 fold higher abundance of these organisms (and their CRISPR arrays) in the ecosystems than other microbes (Figure S2, [7, 21, 45]). Additionally, two versions of a CRISPR DR sequence assigned to *Ca.* Altiarchaea were highly conserved across these ecosystems (Figure S3, Table S4). DR sequence 1 occurred in all four ecosystems, whereas DR sequence 2 was only found at the HURL and the ACLF site (Table S4). While all DR sequences from the four sites were previously unknown in the CRISPRmap database, DR sequence 1 in orientation 1 and DR sequence 2 in both orientations structurally resembled motif 13 and 12 of the database, respectively. All four sequences form thermodynamically favorable secondary structures and carry an AAA(N) motif (Figure S3), indicating that both strands of the CRISPR array could theoretically be transcribed.

### Eight novel viral clades with genome relatedness across continents show infection histories with Altiarchaeota

Using matches of Altiarchaeota spacers to protospacers, we were able to identify 13 predicted viral genomes in three out of the four sampling sites (Table 1). No viruses targeted by *Ca.* Altiarchaea spacers could be predicted for GA and the HURL site at 140 m depth. Only two out of the 13 predicted viral genomes, i.e., the 20.8 kb Altivir_2_MSI_BF_2012 and the 22.6 kb Altivir_8_HURL, had hits in the VOG database (Table S5), carried viral hallmark genes and were circular, prompting us to classify them as viruses. The others were designated as putative viruses according to our classification scheme (Figure S1). All 13 were categorized as lytic viruses according to VirSorter [82]. Four predicted viruses were circular and thus complete in their genome sequence (Figure 1, Table 1). The 13 viral genomes formed eight monophyletic clades based on the VICTOR analysis, representing potentially eight individual genera (VICTOR threshold for genus was 15.8% nucleotide based intergenomic similarity). Viral genomes in the Altivir_1_MSI and Altivir_2_MSI clades were recovered in all sampled time points from the MSI site (Table 1, Figure 2), both in the cellular and virus enriched fractions. The Altivir_2_MSI genomes recovered from the virus enriched fraction were fragmented and thus excluded from further analysis. Intergenomic pairwise similarities for the four Altivir_1_MSI varied between 99.3-99.7%, this clade representing a single viral species (the threshold for species demarcation was 95% similarity). The three Altivir_2_MSI genomes had similarities between 87.0-96.1% and represented two viral species (Figure S5). Using vConTACT2, Altivir_1_MSI formed a cluster with Altivir_6_ACLF, a putative virus from ACLF, with whom it shared five protein clusters (Figure 2, Figure S4). This relatedness between viruses from highly distant subsurface ecosystems was further supported by the VIRIDIC analysis, which showed an intergenomic similarity of 11.2% for the Altivir_1_MSI/Altivir_6_ACLF pair (Figure S5), and by the VICTOR analysis, which placed them in the same monophyletic clade (Figure 2). All remaining viruses apart from Altivir_1_MSI, Altivir_2_MSI, and Altivir_6_ACLF were designated by vConTACT2 as unclustered singletons. Only one protein cluster was shared between Altivir_2_MSI, Altivir_4_ACLF, and Altivir_8_HURL, and no protein clusters between the remaining 3 viruses, indicating that all these viruses are distant from each other (Figure 2). In total, these eight viral genera were affiliated to seven vConTACT viral clusters (Altivir_1_MSI/Altivir_6_ACLF grouped together), clusters unrelated with previously published viral genomes in the RefSeq94 database (Figure S4). Comparing the genomes of Altivir_1 and Altivir_2 individually across the samples from 2012 and 2018, we identified different developments of the two viral genomes. Altivir_2_MSI clade presented gene content variations between the genomes from different samples (Figure S6), which is in agreement with the fact that it represents two viral species. By contrast, Altivir_1_MSI accumulated multiple single nucleotide polymorphisms and only represented strain level variations of the same species (SNPs) (Figure S7, Table S6).

**Figure 1:**
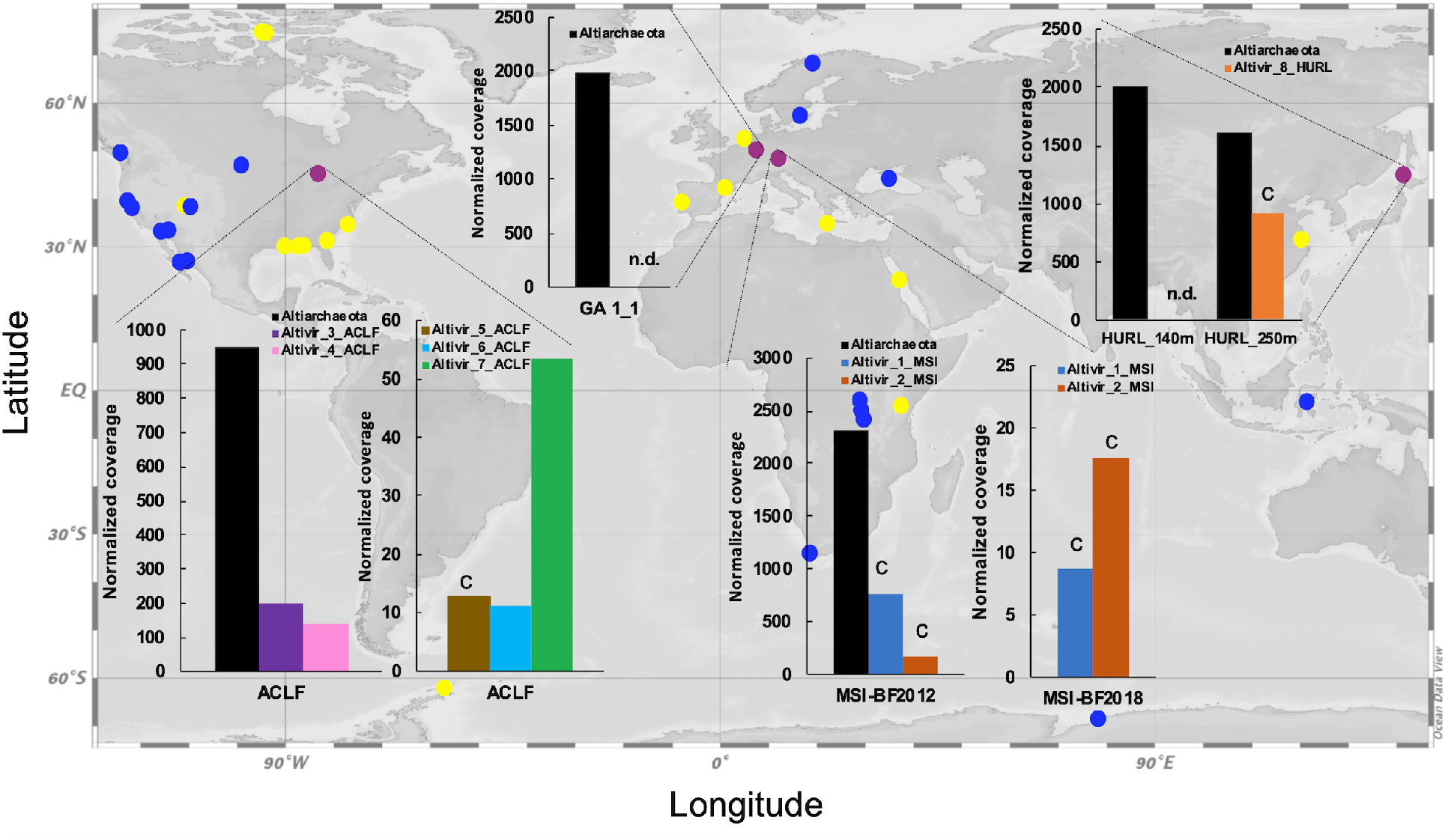
Global distribution of abundant Altiarchaeota (group Alti-1) and their predicted viruses in Altiarchaeota hot spots. Distribution analysis is based on 16S rRNA gene sequencing (yellow dots) and the detection of *hamus* genes in metagenomes from IMG (blue and purple dots). Purple dots correspond to the four investigated sites of this study. Normalized host and virus coverage are given for the four subsurface habitats: Alpena County Library Fountain (ACLF, MI, USA), Horonobe Underground Research Laboratory (HURL, Japan), Mühlbacher Schwefelquelle Isling (MSI, Germany) and Geyser Andernach (GA, Germany). Percent relative abundance of dominant Altiarchaeota compared to other community members is shown in Table S1. Only Altivir_1_MSI and Altivir_2_MSI obtained from biofilm (BF) samples are shown. The letter “C” indicates circular genomes. n.d.: none detected. World map has been generated using Ocean Data View[99].

**Figure 2:**
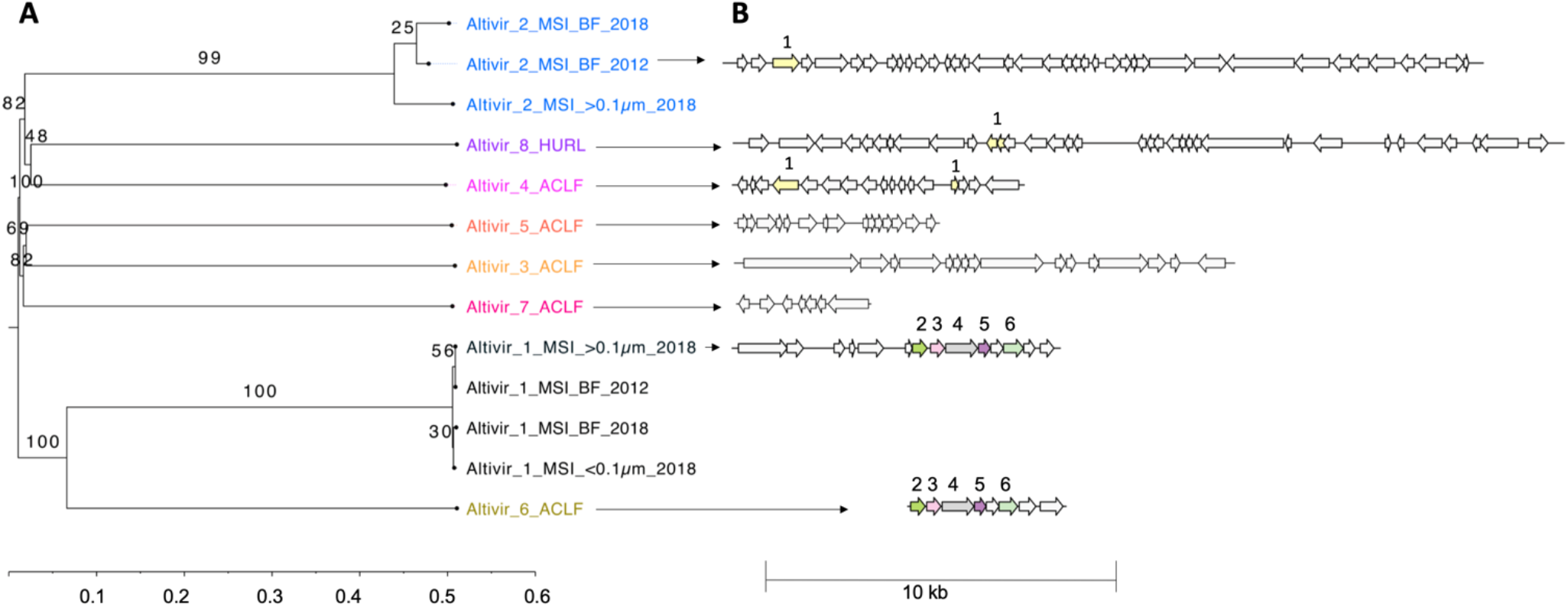
Phylogenomic Genome-BLAST Distance Phylogeny (GBDP) tree of Altiarchaeota viruses and viral proteins clusters. A) The tree was inferred using the distance formula D0 yielding average support of 69%. The numbers above branches are GBDP pseudo-bootstrap support values from 100 replications. The branch lengths of the resulting VICTOR [65] trees are scaled in terms of the respective distance formula used. The tree shows that the eight predicted viral genomes were assigned to the same family, to eight different genera and ten species (which is the result of having multiple genomes included for Altivir_1_MSI and Altivir_2_MSI). B) Protein clustering across all viruses revealed that Altivir_1_MSI_>0.1μm_2018 and Altivir_6_ ACLF, originating from different continents, have five protein clusters (2-6) in common. Altivir_2_MSI_BF_2012, Altivir_4_ACLF and Altivir_8_HURL shared only one protein cluster (1). Open reading frames of the viral genomes were predicted using MetaGeneAnnotator [100] and further translated with the R package seqinr v.3.6.-1.

Host-virus ratios based on metagenome read mapping varied greatly (between 1.8 and 301.9; Table 1) with the smallest ratios of 1.8 for Altivir_8_HURL, followed by 2.3 for Altivir_1_ MSI_>0.1μm_2018, 2.8 for Altivir_2_MSI_>0.1μm_2018 and 3.0 for Altivir_1_MSI_BF_2012. Abundance of viruses in the planktonic fraction (>0.1μm) suggests profound concentration of Altiarchaeum BF on the 0.1μm filter membrane during the filtration process. Relative abundance of viruses based on normalized coverage ranged between 9 (Altivir_1_MSI_BF_2018) and 913 (Altivir_8_HURL, Figure 1, Table 1). The genomes of Altivir_2_MSI_ BF_2018 and Altivir_8_HURL carried short CRISPR arrays with one spacer each, but spacers from these mini-CRISPR arrays did not match other viruses in the respective ecosystems (data not shown).

Several predicted viruses carried genes with matches in public databases, which included methyltransferases, tetratricopeptide repeat proteins and proteins of the AAA family ATPase (Table 1, Table S7). We found the circular Altivir_8_HURL genome to have 36 genes including the auxiliary metabolic gene (AMG) phosphoadenosine phosphosulfate reductase (PHMMR, e value: 6.1e-48) probably facilitating assimilatory sulfate reduction in the host. Altivir_1_MSI was of particular interest for this study, as it was highly abundant in the metagenome MSI_BF-2012, recruited many spacer hits and only five out of its 14 proteins could be annotated (Table 1, Table S7). These included a nuclease (HHpred, e value: 2.8e-19) and a Mucin-like domain-containing protein (PHMMR, e value: 4.2e-23) having a likely role for attachment to the host cell surface. In sum, only 16.4% of the 159 proteins across all Altivir genomes have a putative function assigned rendering the remaining genes of yet unknown function as genetic dark matter (summary of annotations in Table 1).

### Genome-informed microscopy reveals a lytic lifestyle for Altivir_1_MSI

We selected the *in silico* predicted Altivir_1_MSI viral clade for visualization by genome-informed microscopy, due to its high abundance at the MSI site, and despite the lack of viral hallmark genes. Virus-targeted genomeFISH with a probe containing eleven double stranded polynucleotides was successfully implemented to visualize the distribution of the circular genome of Altivir_1_MSI within altiarchaeal BF (Figure 3A, Figure S8). In contrast to the negative control with a non-matching probe (Figure S9), our target probes enabled us to detect altiarchaeal cells containing Altivir_1_MSI. Multiple cells were surrounded by halo signals, corresponding to a viral burst [83] and providing evidence for Altivir_1_MSI being an active virus and lysing Altiarchaeota cells.

**Figure 3:**
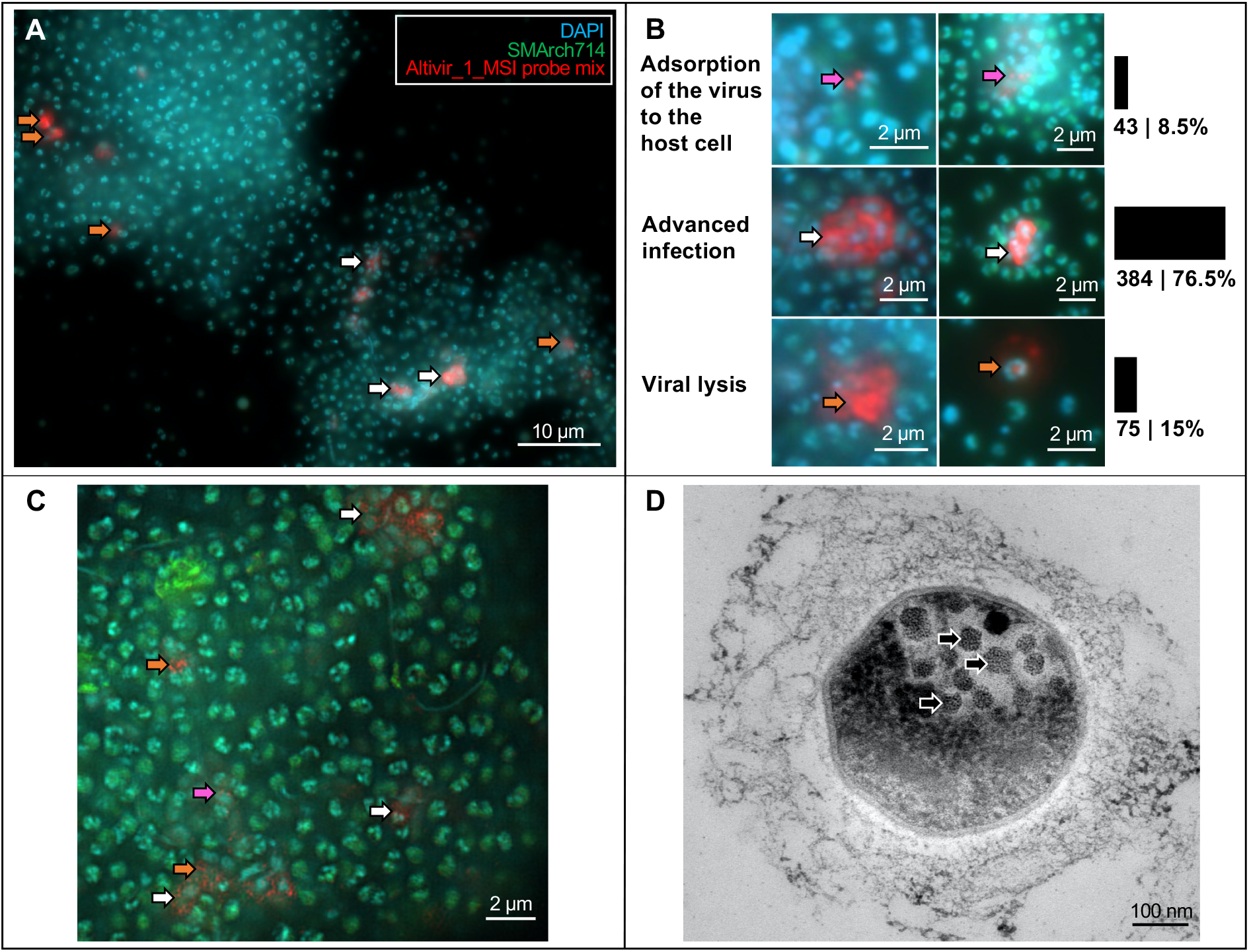
Visualization and quantification of Altivir_1_MSI-infected and non-infected Altiarchaeota biofilm (BF) cells from MSI site. For all virus-targeted genomeFISH experiments shown here, an Altivir_1_MSI probe was used for detecting viral infections. BF material was visualized with filter sets for DAPI (blue, cells), ATTO 488 (green, 16S rRNA signal) and Alexa 594 (red, viral genomes), and merged for analysis and display purposes. A) Virus-targeted genomeFISH displays the interactions between Altiarchaeota cells and their virus shown as red dots. For unmerged imaging data, see Figure S8. B) Different infection stages with Altivir_1_MSI. The enumeration performed with a regular epifluorescence microscope was based on 18 411 archaeal cells and categorized into three infection stages. Purple arrows indicate exemplary viruses that attach to host’s cell surface, white arrows show advanced infections and orange arrows bursting cells with free viruses. C) Coupling virus-targeted genomeFISH with structured illumination microscopy showed extracellular signals of tiny fluorescently labeled viral particles attaching to Altiarchaeota’s cell surface but also in a free state as presumably released virions. D) Transmission electron microscopy revealed intracellular virus-like particles.

A total of 18 411 altiarchaeal cells and 502 viral infections (co-localization of Altivir_1_MSI and *Ca.* Altiarchaea signals) across 18 samples/BF flocks were analyzed via fluorescence microscopy and categorized into three main infection stages: i) viral adsorption to the host cells (8.5%), ii) advanced infection with intracellular virus signals and ring-like signals around the cells (76.5%), and iii) cell lysis with bursting cells and release of virions (15%) (Figure 3B). Super-resolution microscopy further showed extracellular signals of small fluorescently labeled particles, which we interpret as individual viral particles (Figure 3C). This observation was further supported by ultra-thin sectioning and transmission electron microscopy, which revealed many intracellular virus-like particles associated with *Ca.* Altiarchaea cells. These particles had an average diameter of 50 nm (SD ± 7 nm), as measured across eleven host cells (Figure 3D). The high percentage of cell lysis associated with the virus signals along with the high abundance of the virus in the planktonic and viral fraction suggests a lytic lifestyle for Altivir_1_MSI.

### Spatio-temporal heterogeneity of predicted infections and CRISPR-Cas mediated immunity of *Ca.* Altiarchaea

To investigate the development of virus immunities over time, we compared the publicly available metagenome from MSI (taken in 2012) to a newly sequenced BF sample from 2018. We also analyzed the planktonic microbiome (>0.1 μm) and a viral fraction (<0.1 μm), i.e., after 0.1 μm filtration and FeCl_3_ precipitation. We compared the change in relative abundance (normalized coverages) of *Ca.* Altiarchaeum, Altivir_1_MSI, Altivir_2_MSI and *Ca.* Altiarchaeum CRISPR spacers across these samples (Figure 4) with *Ca.* Altiarchaeum being the most dominant microbe in each sample (Figure S2). While the planktonic microbiome showed a tremendous diversity based on *rpS3* sequences (238 different organisms, Figure 4), the diversity was quite restricted with 19 and 17 organisms in MSI_BF_2012 and MSI_BF_2018, respectively. Please note that there is a difference between the number of organisms detected in the rank abundance curves (Figure S2) and those reported in Figure 4 as the latter were normalized to read abundance to ensure comparability.

**Figure 4:**
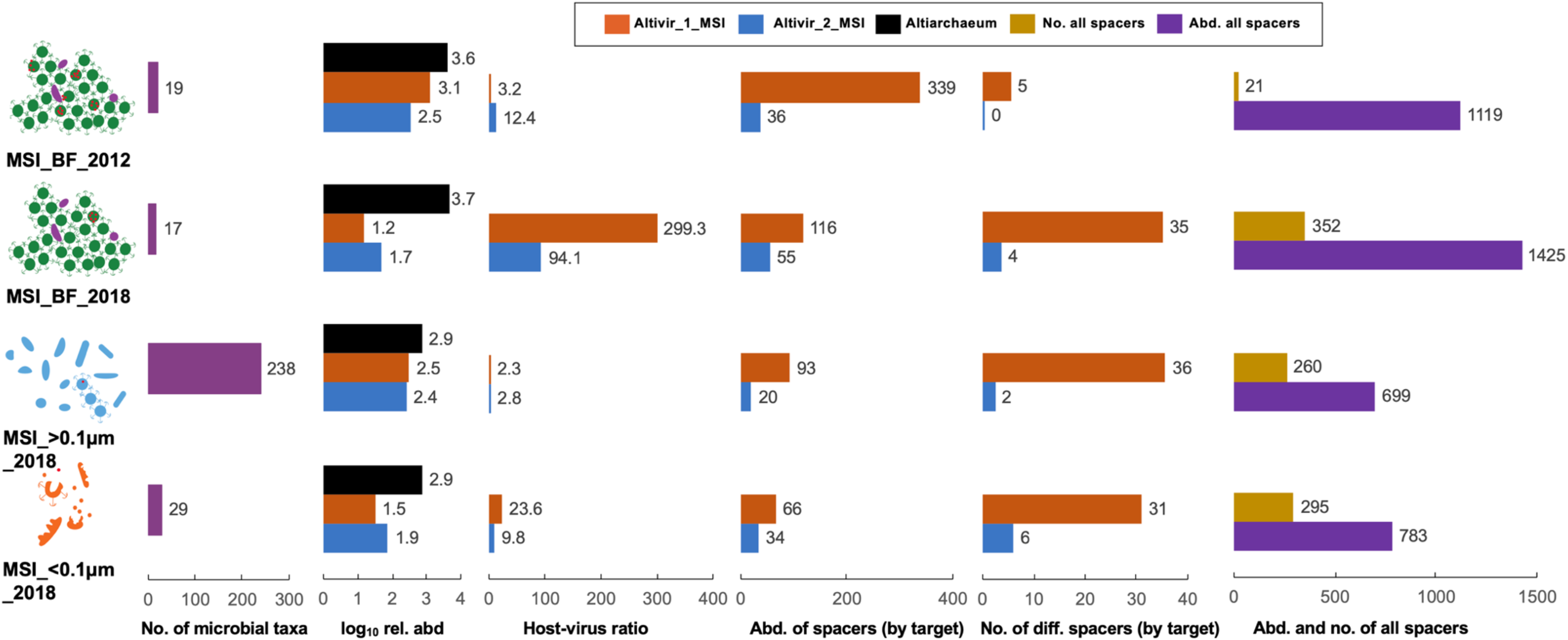
Development of host-virus and spacer dynamics from 2012 to 2018 based on metagenomics. Predicted viruses became less abundant from 2012 to 2018. Considering the biofilms (BF), total spacer abundance and numbers increased from 2012 to 2018 while those matching Altivir_1_MSI decreased in abundance but increased in numbers. Number (no.) of microbial taxa refers to the number of different prokaryotes in a sample detected via *rpS3* rank abundance curves (normalized by sequencing depth). Host-virus ratio is calculated from host and virus coverage based on read mapping. Abundance and number of different spacers were normalized to minimum relative abundance (rel. abd.) of the host based on read mapping;

Both predicted viral genomes (Altivir_1_MSI_BF_2012, Altivir_2_MSI_BF_2012) declined in abundance when comparing the BF sample from 2018 to the sample from 2012, and at the same time the host:virus ratio increased (Figure 4). While Altivir_1_MSI was more abundant in 2012 (host-virus ratio=3.2) compared to Altivir_2_MSI relative to its host (host-virus ratio=12.4), the pattern reversed in 2018 for the BF sample (host-virus ratio=299.3 compared to 94.1 for Altivir_1_MSI and Altivir_2_MSI, respectively). Both, the total spacer abundance and the spacer diversity increased from 2012 to 2018 in BF samples; i.e. from 21 to 352 (number of different spacers) including an ~20% increase in the number of spacer clusters that were singletons in the dataset (Figure S10) and from 1 119 to 1 425 (abundance of spacers). The abundance of spacers matching the genome of Altivir_1_MSI_BF_2012 decreased from 339 to 116 spacers, whereas it increased from 36 to 55 for Altivir_2_MSI_BF_2012 in BF samples from 2012 to 2018. For both targets the number of different matching spacers increased over time, in line with the development of the total spacers in this ecosystem (Figure 4). Because planktonic *Ca.* Altiarchaeum cells (diameter: 0.4-0.6 μm) cannot pass the 0.1 μm pore-size filters, it is more likely that lysed *Ca.* Altiarchaeum cells ended up in the <0.1 μm fraction in 2018 allowing binning of their genomes including the CRISPR system with low complexity of spacers from this fraction. The MSI_>0.1μm_2018 fraction and the MSI_<0.1μm_2018 contained about half the number of total spacers of the MSI_BF_2018 sample, although the number of different spacers displayed less variability. Spacers from these samples hitting the viral targets were often reduced in abundance compared to BF-derived CRISPR spacers (Figure 4).

Congruent with the decline in relative abundance of Altivir_1_MSI based on metagenomic analysis, we also observed heterogeneous infections of BF flocks via imaging. Some BF showed no infection with Altivir_1_MSI at all (Figure 5A, Figure S11), which aligns well with the decrease of the virus in the metagenomic data of the BF from 2012 to 2018. By contrast, we observed very few BF flocks that showed an extremely high infection and accumulation of rod-shaped microorganisms (Figure 5B, Figure S12&S13). The observed heterogeneity of infections in BF supports the aforementioned heterogeneity related to CRISPR resistances against Altivir_1_MSI with high spacer diversity dominated by singletons.

**Figure 5:**
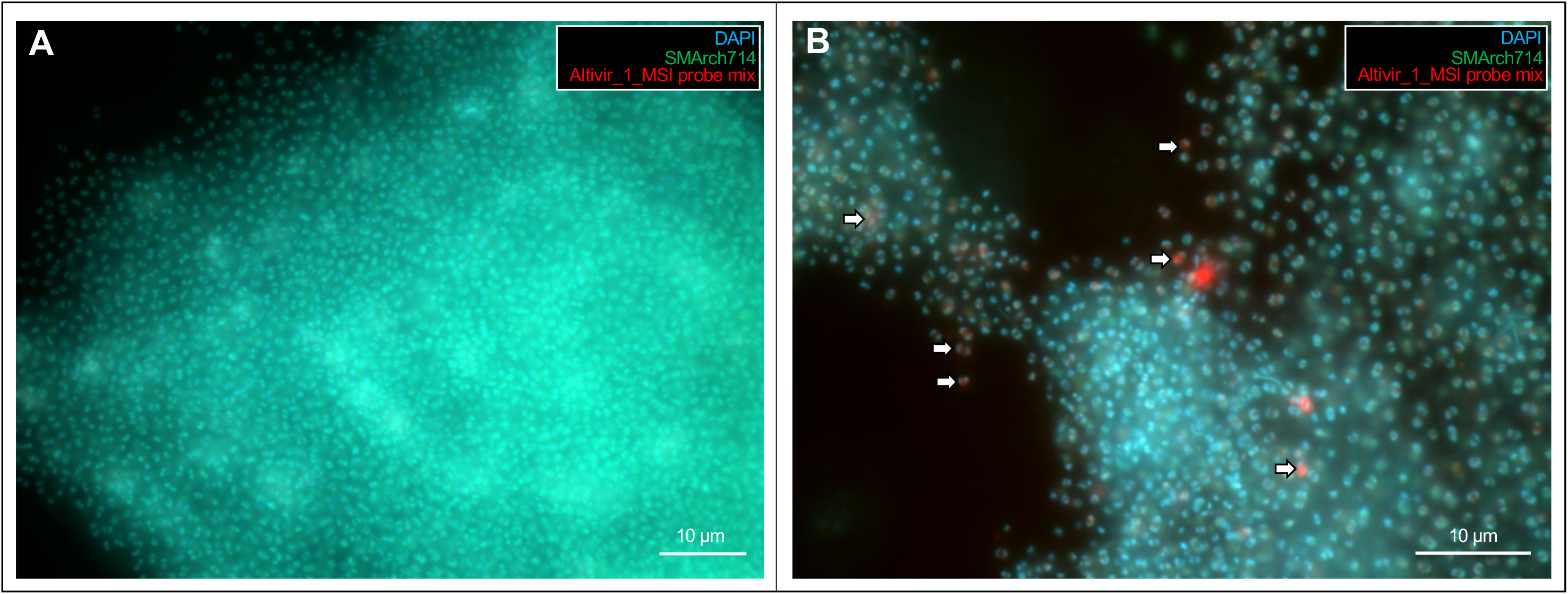
Virus-targeted genomeFISH of two individual Altiarchaeota biofilm (BF) flocks depicting (A) a dense BF flock without infections and (B) one highly infected BF flock. For all virus-targeted genomeFISH experiments, an Altivir_1_MSI probe was used for detecting viral infections. White arrows indicate exemplary virus-Altiarchaeota interactions (advanced infections). BF material was analyzed with filter sets for DAPI (blue, cells), ATTO 488 (green, 16S rRNA signal) and Alexa 594 (red, viral genomes), and then the different fluorescent channels were merged for analysis and display purposes. For unmerged imaging data, see Figures S11&S12.

## DISCUSSION

Although life in the deep subsurface contributes significantly to overall biomass and microbial biodiversity on our planet, its low accessibility leaves host-virus interactions—especially those of uncultivated hosts—highly enigmatic. Detection of novel and uncultivated viruses missing conserved sets of hallmark genes has been an ongoing challenge in viromics [84]. Using a combination of bioinformatics and virus-targeted genomeFISH, we were able to visualize infections of Altiarchaeota with a hitherto unknown, presumably lytic virus from a subsurface ecosystem. We expanded the diversity of this virus detected in Germany (MSI) by identifying a distantly related viral genome also infecting *Ca.* Altiarchaea in North America (ACLF site). This suggests genomic relatedness of archaeal viruses over long distances, a phenomenon known for bacteriophages [85, 86] as well as a common infection mechanism. While Altiarchaeota viruses do not represent an exception regarding the high degree of protein annotations that remain dark matter, some proteins such as modification methylases or the phosphoadenosine phosphosulfate reductase were previously also detected on a *Methanosarcina* and other archaeal virus genomes [87, 88]. Altiarchaeota can apparently employ two different CRISPR systems (Type I and III) with high similarity of their DR sequences across larger geographic distances; although the reasoning behind two independent CRISPR systems remains unclear, an additional Type III system might mediate resistance against plasmids carrying matching protospacers but lacking a protospacer-adjacent motif [89], or against viruses that overcome Type I systems as previously described for *Marinomonas mediterranea* [90] and *Sulfolobus islandicus* [91]. Indeed, archaeal viruses found ways to interfere with CRISPR Type III systems by using a plethora of anti-CRISPR proteins [92, 93]. However, we could not detect homologs of these proteins in viruses targeting *Ca.* Altiarchaeum.

Based on the metagenomic data collected in 2012 and 2018, it appears that the host prevailed in the arms race between Altiarchaeaota and the visualized virus (Altivir_1_MSI). The number of different spacers matching this virus increased towards 2018 with a prominence of spacer singletons in the metagenome (Figure S10), while the actual abundance of spacers matching Altivir_1_MSI decreased as did the abundance of the viral genome itself. CRISPR spacer diversification might be a successful response to an increasing number of variants in the genome of Altivir_1_MSI (Figure S7) suggesting its mutations. As a trade-off, spacer diversification might allow the host to decrease the total abundance of spacers. We conclude that CRISPR spacers were diversifying to mediate a greater bandwidth of resistances against the virus, which worked in favor for the overall population of Altiarchaeota in this ecosystem. Moreover, the diversity of singleton spacers indicates a very heterogeneous Altiarchaeota population, which might be related to heterogeneous infections of the BF as visualized via virus-targeted genomeFISH.

Our dual approach of coupling metagenomics to fluorescence microscopy enabled us to follow precisely the terrestrial subsurface predator-prey relationship of *Ca.* Altiarchaeum and one of its viruses. Our data suggests that the novel identified virus is lytic and challenge the current paradigm that lysogeny prevails in the subsurface [29, 41]. Instead, our data indicate that the “kill-the-winner” theorem–lytic viruses targeting abundant ecosystem key players [94]––also strongly applies to subsurface ecosystems. We currently do not know if this statement can be transferred to other subsurface ecosystems that have low cell counts [2] as viruses might struggle with finding a new host. However, replication measures of bacteria in the ecosystem with the visualized virus suggest that microbial proliferation is similar to other oligotrophic systems [46]. Lytic infections in subsurface microbial hosts might thus launch heterotrophic carbon cycling similar to the viral shunt in the marine environment [95]. In fact, recent lipidomic analyses coupled to mass-balance calculations provide evidence that subsurface environments dominated by Altiarchaeota are completely fueled by these organisms’ carbon fixation, transferring organic carbon to heterotrophs in the community [96]. This process might be the basis for microbial loops [97] as we see the accumulation of rod-shaped microbes around Altiarchaeota when they are lysing due to viral attacks (Figure S13).

Here, we provide underpinning evidence for the frequent viral infection of a globally abundant, autotrophic key player of the subsurface carbon cycle. We discovered that one virus with a lytic lifestyle was even capable of infecting host cells in a dense BF, which is known to provide some protection from viral infection [98]. Subsurface ecosystems such as aquifers remain understudied regarding host-virus dynamics because of limited access, low microbial biomass and limited cultivation success. Our results presented here provide an experimental proof of concept for an ongoing host-virus arms race in the continental subsurface characterized by constant viral infections and cell lysis of subsurface microbes, followed by their own and their viruses’ diversification.

## ACKNOWLEDGEMENTS

This work received funding by the Alfred P. Sloan foundation (grant number G-2017-9955), the Ministry of Culture and Science of North Rhine-Westphalia (Nachwuchsgruppe “Dr. Alexander Probst”), and the NOVAC project of the German Science Foundation (grant number DFG PR1603/2-1). We acknowledge sequencing of MSI metagenomes within the Census of Deep Life Sequencing call 2018, phase 14 project “Development of novel archaeal viruses and the corresponding CRISPR arrays of a highly abundant carbon fixer in Earth’s crust”. We highly appreciate the technical assistance of Jannis Becker for protein annotations of viral scaffolds, and Sabrina Eisfeld for laboratory maintenance. The work conducted by the U.S. Department of Energy Joint Genome Institute, a DOE Office of Science User Facility, is supported under Contract No. DE-AC02-05CH11231. Furthermore, we are also grateful to the Imaging Center Essen (IMCES) under the direction of Matthias Gunzer for access to an inverted epifluorescence microscope with structured illumination and for the respective training carried out by Alexandra Brenzel.

## COMPETING INTEREST

The authors declare no conflict of interest.

## Notes

### Competing Interest Statement

The authors have declared no competing interest.

