## Supplementary Information for "Genome-informed microscopy reveals infections of uncultivated carbon-fixing archaea by lytic viruses in Earth’s crust"

[2]DOE Joint Genome Institute, 2800 Mitchell Drive, Walnut Creek, CA, 94598, USA

Content:

1. Supplementary Material and Methods
2. Supplementary Discussion
3. Supplementary Figures
4. Supplementary Tables
5. References

**Table S1: List of accession numbers for metagenomic reads, assemblies and genomes deposited in public databases.** ACLF=Alpena County Library Fountain, ENA=European Nucleotide Archive, GA=Geyser Andernach, HURL=Horonobe Underground Research Laboratory, MSI=Mühlbacher Schwefelquelle Isling, SRA=Sequence Read Archive

**Metagenomes**

| Sample | Database | Bioproject accession | Biosample/Run accession | Release date |
|---|---|---|---|---|
| MSI_BF_2012 | ENA | PRJEB6121 | ERR628383 | released |
| MSI_BF_2018 | SRA | PRJNA628506 | SRR11614987 | 03.05.21 |
| MSI_>0.1µm_2018 | SRA | PRJNA628506 | SRR11614986 | 03.05.21 |
| MSI_<0.1µm_2018 | SRA | PRJNA628506 | SRR11614988 | 03.05.21 |
| HURL_250m | SRA | PRJNA321556 | SRR3546457 | released |
| HURL_140m | SRA | PRJNA321556 | SRR3546456 | released |
| GA_1_1 | SRA | PRJNA627655 | SRR11600163 | 03.05.21 |
| GA_1_2 | SRA | PRJNA627655 | SRR11600162 | 03.05.21 |
| GA_2_1 | SRA | PRJNA627655 | SRR11600161 | 03.05.21 |
| ACLF | SRA | PRJNA340050 | SRR4293692 | released |

**Assemblies**

| Sample | Database | Bioproject accession | Biosample/Run accession | Release date | % Rel abund. Altiarchaeota* |
|---|---|---|---|---|---|
| MSI_BF_2012 | ENA | to be submitted | | | 90.7 |
| MSI_BF_2018 | ENA | to be submitted | | | 97.6 |
| MSI_>0.1µm_2018 | ENA | to be submitted | | | 38.5 |
| MSI_<0.1µm_2018 | ENA | to be submitted | | | 74.9 |
| HURL_250m | ENA | to be submitted | | | 68.9# |
| HURL_140m | ENA | to be submitted | | | 77.1# |
| GA_1_1 | ENA | to be submitted | | | 86.1 |

| | | | | | |
|---|---|---|---|---|---|
| GA_1_2 | ENA | to be submitted | | | 87.1 |
| GA_2_1 | ENA | to be submitted | | | 88.1 |
| ACLF | ENA | to be submitted | | | 52.0 |

**Altiarchaeota Genomes**

| **Sample** | **Database** | **Bioproject accession** | **WGS accession** | **Release date** |
|---|---|---|---|---|
| MSI_BF_2012 | ENA | PRJEB6121 | CCXY01000000 | released |
| MSI_<0.1µm_2018 | SRA | to be submitted | | |
| HURL_250m | SRA | to be submitted | | |
| HURL_140m | SRA | to be submitted | | |
| GA_1.1 | SRA | to be submitted | | |
| GA_1.2 | SRA | to be submitted | | |
| GA_2.1 | SRA | to be submitted | | |
| ACLF | SRA | to be submitted | | |
| Altivir_1_MSI | SRA | to be submitted | | |

*Percent relative abundance compared to other community members based on *rpS3* sequence analysis.

[#] since the metaspades assembly of the Altiarchaeota genome was extremely fragmented and the *rpS3* gene did not assemble we used a scaffold carrying the ribosomal proteins L30 and L15 for coverage estimation of the dominant Altiarchaeum.

**Table S2: Sampling information for biofilms (BF) collected from the Mühlbacher Schwefelquelle, Isling, Germany (MSI)**

| Sample | Time of sampling | Analyses | Figures |
|---|---|---|---|
| BF samples for virus-targeted genomeFISH | 17 - 19.01.2019 | Fluorescence microscopy | Figure 3 (A-C), Figure 5, Fig. S8, S9, S11, S12, S13 |
| BF samples for transmission electron microscopy | 17 - 19.10.2018 | Transmission electron microscopy | Figure 3 (D) |
| BF, >0.1μm and <0.1μm fraction | 17 - 19.10.2018 | Metagenomics | Figure 1, 2, 4 |
| BF | 11.2012 - 04.2012 | Metagenomics | Figure 1, 2, 4 |

**Table S3A: Sequence of the circular viral genome Altivir_1_MSI_BF_2012**. The regions corresponding to polynucleotide binding sites are represented in different colors. Poly=polynucleotide

| | |
|---|---|
| via:<br>>VIRSorter_NODE_1844_length_8923_cov_1978_89-circular-cat_3_modified<br><br>11 polynucleotide probes (each 300 bp) from IDT (Leuven, Belgium):<br><br>Altivir_1_MSI_probe_1<br>Altivir_1_MSI_probe_2<br>Altivir_1_MSI_probe_3<br>Altivir_1_MSI_probe_4<br>Altivir_1_MSI_probe_5<br>Altivir_1_MSI_probe_6<br>Altivir_1_MSI_probe_7<br>Altivir_1_MSI_probe_8<br>Altivir_1_MSI_probe_9<br>Altivir_1_MSI_probe_10<br>Altivir_1_MSI_probe_11 | CCTGCCCAGATACTACATCTCAAACATTCCTCCAGGCATATCTTTGTGCCATACAATATCAA<br>AAAACAGGCGTAAAAGACGAACACGACTTTGAAATTTATTCCAGTTTTGACATTGAAGATCA<br>AAACGAATATGTTTATATTTCATCTGAAAGAGAAAAGAGTGACTTGTTCGAACAACAAAACG<br>CAAACTGGTATTTGATTGAAAGAGATTTACTTGGAAAAATCACGTACATCGACACAGAAACC<br>GGAGATGATTTTTCATTCAAAGAAATTGAGTCTGGGAAGCGAAAAATAGTCGAAGCGGGTGC<br>TGGAAGCATCTTCAAACTTCTCGGAAAACTAAAAACAAAATTTGGTAATTTTTTGCTCGCCA<br>ACGCAAGCCAAAAATCACTGTGGATATTAAAAAAAGAAAAATCATCAAGCTGGACAAAAAAA<br>CGGGAGCGTTCGTAATTGCCACAGGTATTTCTTCCACAACACTCATTACATCCCTCTTAAGC<br>CAAACCTCATCAACACTTCTCGCAAAAGTTCGCCTTCTCATTGAAATGCAGGGAATAGCACT<br>TTTCACGGATTTCATTGCTGAAGAAGCAGTTCAAAGTGCAGGAATGGGTGTCTTCATAGCGA<br>AAGGTCGCGACACAAAAACACTCGAACATGCTATCTCTACATACAAAAATATATATTCTCTG<br>GCTGAAAATGTACATCAATACGCACAGTGGAATATAGCGAAAAATGCCTTTGATGCCTTTTT<br>CCTCTCCGCAAAAGAAAACATAGTGATATATGAGGCACTCCTTGAAAATGCGAAAGAAACAG<br>CTGCACTTCGCGGAGAAACAGAGGACGCGGGAAAAGTCACTATATATTCTACCGAAAAATAT<br>GCAATATACCTCAACGGAATTAATACAAAAAAATACTCTCCGTCAGTCATACGAATTCCACC<br>AGGAACGCATACGATAGACCTGAAAATGAAAGACAAAATCACATATACAGAAACATTTTCAA<br>TCGAAAAATATGGCATCAAAAACATCGGGTCTGTTTCAAAAAAAATTGAACTACAAAACACA<br>ACAAAATGTCTTGATGTTGCCAACGGAACAACGATAATTCTCAACGATTTTCAACAAATCGT<br>ATTGGCAGGCGTAAAACAAAAAATTTTAACAGCAGCAGCTGAAAAAGCATCAAAAAACTGGC<br>TTCGTAAAACATGCGAAACAAGGAATATTCGCATCGAAACCGATACAGAAACTCATAACGAA<br>CATGGACAATTGTTAGGCTGGATTTTTCTTGGGGATCTAAACGTAAACGTTGAATCCGTAAA<br>ACGCGGATTTACAATAAGTAACAAAACAACAAAAAAATATGAAGATGAAATAAAAGAAGCGG<br>AAAAATTTGCCGAAAAATAATAAATTACAACAAAAAATGTTCGGAAGAGCAAATTTTTTGCC<br>AAAACACCAGAACAAAAAAGACCTACTATGGAAAGACATAGATGACTTTGTCAATTTTTATT<br>TCTTGAATCAAAAAATTTCAAAATCATACGAATTTGAACTTTACAACCCACGATGGTTTAAC<br>TCCACAAAAGCTTGGCAAATTTCATGCGCAAAGAAAATAGACCTGGTTGTAAATCGTAACGA<br>TGAAATTTATATTTTTGAAACCGACCCCCAAATTCTCGATGAAACAATCGGGCAACTTCTCA<br>CCTATCGTTTTTGGTATGAAAAGCAGGAACAAAAAAAAGTGAAAAAAATGATAGCACTATGT<br>GGGGAAATCCAAGAGCAAACTGCCGAAGTAGCACTTCTGTATGGAATAGAAGTCTGGAAAAC<br>AAATTTTCCTTCTGGATATACTCAGTATGTTTCTATGTACTAAATATGTTTCTATCTATTGA<br>ATTCATCGAAATTAAATTTTTTCTTGCCTAAATTACCCTCAAAAGTATGGGGAGCGACAAAT<br>AGAAGATTGTCCTCTTGGCCAATCTTCTAGGGATATTAAATTTTCCCCCGCCTAAATTGTTC<br>TTAAAAGTACGGGGAGCGACCGGGGGGGCTCAAACAAAAAAAAAAATGTGAATTTTGATTTTT<br>TTTTTGTTTGCTCGGAGCCTGGCGGGCTCTTGCCCGCCCGGCTCCGAGGTGGAGTCTGGTTT<br>GGGGGCAAGCGGCTTCGCCGAAAATTCTTTGGAATTTTCCCCCAAACCATCCTCCACCTCTG<br>AGCCTATCGGCTCCGAGACGCCCCCCAGCCTATCAAAACATCAGAAAAATTCTTGGGCTACG<br>GGTATGAGCCCTACGCCCAAGAATTTTTCAGAGGTTTTGATAGGCTATATATCTTCTCATTG<br>TTATAAAAATTCTCCACAAAGTCTTCCTAAATTTTCCTAAATTTTCCTAAATTTTCCTAAAT<br>TTTTCCCTCCCTTGCTCCCTCATAAGACCGCTACCATCAAACAAACTTAAGGAATACTATAT<br>ATGCCACCTCATAAGTATCTTTTTTCTGCGAAGAAAAAAGATACTTAAAAGGGCTCTGGCAT<br>ATATATCCAACCATAAGTTCGTTTGAAGGTGCGAGCTTATGAGGGGCAAGGGGGGACGAATA<br>CAAATTTTCCTAAATTTTTCCGGAAGGTGAACATCGGAGGGGAGTTAAGTCCGAATATAAAT<br>CTCCTACTTGCGAGACAAACGCCAAATTTTCGGAATGGCGAACTCGGAAAGGGGAGTTAAGC<br>TAAATTTCCCACTGCGAGACAAACGCCAAATTTTCGGAATGGCGAACTCGGAAAGGGGAGTT<br>AAGCTAAATTTCCCACTGCGAGACAAACGCCAAATTTTAAAATTTCAAAATAATAAACAAAC<br>GACCATGAAAAATATAAATCCCACAAATGTGGTGATAGGACGCAAAAAAAAAATTTGAAGAAG<br>AAATGAAAGAAAAAAATAGAAGAATATGTGAGGAACCATCAAGAAGAAATAGTAGAGAAAGAA<br>AAAGAGAACATGAATCCGCCTGGAACAGCAGTCGTCTGGCAAGAGTCGGGTGGGATATTTCC<br>CTCTCGCCGGAAAAATCCGAGGGCGATATGACTTGTGATTGCTTCTTCGTTGAGATTAACAA<br>GAAAACACAAACACACACAATATTTTTTATTTTTTATTTTTTAAAAGTTTTGCTCATTCCCG<br>TATTTTTAAACCTTTCACCTTTATTCAAAGGTAAATCTTTTAAATATGGTTATGTGTGCTAA<br>CTTCTGCGGTTTAAACTTTTTAACTAACACACACAAAAAAAAAGTTTATAAAAGAAAGGTAG<br>CCAAAGAAAATTATCAAAAGCAACAAAGTGGTATGTCATCGCTTAACAATTATGAAAGCGAG<br>GTTAAGTATTAATTAATTTTAAACTTTTTATCTTTATTCTTTGTTAGAAAAATTTAATTATT<br>ATTATTATTAAAAAAAATATAAAAAAAAATAAATAAAAAACAATGGAAAATATTTTTGAAAC<br>CGTGATGAAGGAACCAATAGAAATTAAAAAAAAACGCGAACAAATAGCAAGAATAAAATATC<br>TCAAAGACCGACATGGCGTCATTTCCCTATGGCGCGAAGAACCAGTCCAGATAAATCTTCGG<br>GGATATCTCCTGACTATCAGGCGTATCGAAGGTGAAACAGGAACGACAATTTCTGATGTCGC<br>GAAAGAAACGGGAAAAAACATTAAAACCGTCACGAATATTTTTAGAAATCTTGAAAAAATTG |

AAGTGATAAAAATTTCCGAACATGCACATTTCTCTCGAAGAAAAAAAAATAAGAAAAACATA
AATTTTTTCAAATCAAAATTCAAATTGACAACATACATTGAGGAATATTACTCGAATCATAG
TTTTTTCCGAAATCAATATACGTATAGCGATCTCAAACCTGCTGTGATTGAAGCAAAAGAAA
ATGAAAGAGTATCAAAACGCATGGATCTTGAGGAATCTCGTTATTTATGCTCAAAATCTATT
GTGCAGTTTGACCGTGTCTCGCCGCCCTCGCCGGAAAAATCCGGAGGTGATGTTTTACCCTC
ACGGAAAAATCCGGAGGTGATGTTTTACCCTCACCGGAAAAATCCGGAGGTGATGTCTCACC
CTCACCGGAAAAATCCGGAGGTGATGTCTCACCCTCACCGGAAAAATCTCACTGGCGGGATT
TGGATGAGGTATTTCCATCTCGTAAGAAAAATCCGAAGGCGTTATGACCTTCACCAGAAAAA
TCCGAAGACATTTGAATATGACGAATATAAAATATAGTTTTTTTAGGTAACCTTTTTTCCCA
AAAAAGGTTTTATTTAATACAAGAAGTGCACAATAAATACAGTTTTTTCAAACTCTCCTTGT
TAAAAGTTTAATCATACACGATTAAACCTTTGACAAGGAGCATTTTTAAAAACCACCTTCAA
ACGGGGGGGTTACTTCTTGTTAAGGCATTACAGGAATTTTTTCCTCGCCAAAAAAATTTCCG
GACGCAAAATGAAATTTTTCCGGACGCAAGGTGAATCTATATGTATTTTTCTTAAATTTTTT
TTCAAGTAAAAAATATAAAATGACGAGGAATTGACACTCAACCCTGGTGTGGTTTGAGGGTG
AAGGGGGGTAAAATATAGGTTTTATGCCATTTTATTATTTATTTAATATCAAAATTTAATTA
TTTATTTAATATCAAAATTTAATTATTTATTTAATTATTTATTTAATATCCCTATTTTTTAG
GGATATTTTTTTTTTTAAATCGTACTATGTAGTATAAAAAGATACATAAAAGATGAAAATAAA
ATTCAGAAACAACGGAAACAAAATATCAAATTATGCAGATATAGGAACCATATTGACAATAG
CAGCCAGCACGTTTTTTTCATCTTGTTCCCCACAAAATATTTTTGGCTGCTACATAGCCTTC
ACAATTGTTATTTTTCTAATAATAATAAAATATATTTCAGTAAAAAAAACATAAAAACACAA
AATGACAAAATCAGACTGGATGACCGCCACAGAAATAGAAAACGCCCTCAAAACAGGAATAA
AAAATGCAAAAAATGCATTCGAAAATGGGATAGATAATTGTAGGCGAGACCCCATAAAAACC
GCAATTGCAGCAATAACATCGGGAAAATGGGAAAAAAATTTCAAAGATTCCTCAGAAAAATG
GGAAAAAAATCTTTCAAAAATTTCTCTTGAAGACTGGAAGAAAGCCACAATCGCTGGGGCAT
CCAGATATGCTGACAACGCAGCCGACATAGGATCCGAAGAATGGGGAAAATATTACACTAGT
GCAAAATCAACTATCGAAGCAGCATCGTCAGAATTAATCGCAAGCAAACAAGAAAAAGCAGA
TTTTATCAAATTTTACGAGGCAATGTCCAAATTAAAAAACATAGATTAAAAATAAAGGCCCT
TTTTTTTGAAATTTCATAGCAACATCACGATATTAAATATAAAAGCCAAAATAAATTTAACA
ATAAAAAAAATAAAATGGCAATAAAACCAGAGGATGCAATAAAAAAGTATAAAAATAATATA
GACCAAACAAAATACGAAAAAGGAGTTAATGCGGTCAAAACAAGCGTAACCGCAGCCGCGAT
AAAGCAGAAAGAAAGTTTCATCAAAAACACTCTCGCAGGTAAAGACCGCCTCATAAAAAATT
TGGGCAAAGTTTCAACTGATGTATGGAAAACCAAAACAAAAAGAAGTTTTCAAAACTCGAA
GAAAAAGTTATTCTTGCGATTGACTCGGGAAAATGGAACGCAGCCAAAGTTCTCGCTGCGGG
ACAATCCGCGCACGCAACAGCGAAAAATATGAAAAAAGGATCATATTCTGATTCCTATGATC
GATATCTCGCATCACAAAACGCAATTAAAACCGCATGGAAATAATAAAAAAAATAAATATCA
AAAACACAAAATGACAATAATATCATCAAACAACACCAAATCTATTAATGGGACCGAACTCG
TTCTCGAAGCCGATATAGATAAAAGCATTCTCGTGAAAGGAGTGCACGTGCAAGGCTCAGTA
AACGGAAGGGCAACAGCCTACATTGGAGGCGTCACTGTGGGATACTTTCGAACAAACTCAAC
AATTCTCGGAAACCACCTTGAATACCTACGAAGAAATAATAAAAAAGAAATAAATGTACTAA
ATTTCCTTTTTCAAAACAATGTTTTAAGGGCTACCCTCTCGCACCCGGTAATAAATTCACG
ATAAAAAATTCTTCAGCAGTAAATATGTCGGTAATCTATGATATTTACACAGAAGGCGACCA
AACAGAACTGATGGAAAACGGAAAAAACGGAAAAAGTTTAACATATCTTCACTACGGTCAAA
CCCCAAATTTCATCAACACTACTGGTGACACCCTCATATCTAAATCAAACAATCCAATTGAA
TTCTCGGGTTTTCCATTCACCTCAACCGTACCGCCAAAAAGAAAAATAGAGGTCCTCGGAAT
AGCGTTTTCATCTCGAGGTGCAGACGACGGAATCGCCGCAGCAAATTACATCAAAACCACAT
ATCTTAAGGTTATGAAAAACAGAGAAGTTCTTTTCGACGAAGAAAAAAAAGGTTTCATAGCG
ACAGGATCTCAAGTTTTAACCGCAAATACATTCGACGCCGAAAATGGACTTGACATAGGGGG
AGAAGGGACATCTACGTATCAAAAGAATATTCTTTTATTTGACGAAAAAAATTGTATGTGTCG
CAGGCGAAGAACTAAATTTTATCTGGTCAACAACTGTCGCTGCTATACCTGGAAGGTTTGAA
CCAGAAGAGACAGAAATTATGGTCATCTTAAGAGAAACAAAAGAATAAAAAATTTATAAAAA
ATAAATAAAATGACAATTTTATACAAAACTTTTAGTATTGTTTTAACCGCAGACAACCAAAA
ACATAACGCGACATCAGTCCTTACATGTCCAAAAGGAAAAAAAATACATTTTTCCTTTCTCG
GAATCGCACCATACATCACTGAAAACTGCCACATTCAAGTGTTCAAAGACCAAGAACAACTC
ACCGGAGAAGGAATATACCACGCTATTATCGACGATTACACTCACCGGCTTGATTTTGATGT
TGATCTCAACGAAGGAGAAGCAATCACAATCAAGGGATGGAGTATGACAGCATACACGGTGG
AAGGCGTTTATGCGTATGAAGAATTAATTTAAAAAAATTCAAATGTACAAATATATATGTTT
TGAAAACACACTACTAAAAACAGCTAATTTTAAAACATTATGTGAAATTGAAAAATCATTCG
TGGTTGGAGCGTCGATTAAATTTTGGGAAAACCCCTCTTTTTTTCCCTACGACGCGGTAATA
TCTATCTCAGACATAGGTGCTACAAAAAAATATATACTTGATATATCTTATCCGCATATGTT
CATTGCGAATTCAGAAATGTCTTACTTTGGCAAGTGGTCAAATGACACATTCATAGAATTGA
ACCACCAAATAGATATTGATCACAAACTATTGTTTGAAATTTTGGTCACCAACGATCTTCAA
GTTCAAGGATACGTTTTAATCGAGACATAAAAAATGAAGTTTTTAAACCTATATAAAAAAAT
AAGGAAAAAAACAACCATAAGACAAAATCAAACCCCAAAAACCACCGTACCTGAAACATTCG
AAAAAATATTTCTCGGGATATTTGAAAATTTTGAAAAAATAGGGTGGTCTACAACACATTTG

| | |
|---|---|
| | TATCTCTCAAACATTATTAAACGCACTTTTTCGTATATCCTATACCCAACACTCGACGGGAA<br>AAACGCAGGATTTTTCAAAGAAAAAACAGCTGGTGATATCACAACCCTCCAAACCCAGCCTG<br>GAGGAACGAAATTCTTTCTTGACAACAACCTCGTCCGTAGGACGAAAAGAATAGGTATTTCC<br>GCGTATAATATGTCAGACCTAGTAACATGCGGAGTGCATATCAGAGTCCAGGGACCATTCAT<br>CCAATCACCCATAAATGAAGTAAATCTATTCTACTTCACTGTTGCCCCGGTGACGCTCAAAA<br>CATTTGAATTTGATATTGCGTCGTTTGATGTATATCTTATTTTCGAAGCTGTTGGACCCACA<br>GGAGAATCAATAATTTTCAAAACTTATTATTTACCAATATAAAAATGGCTGATTATTTCGAA<br>TTCACCGTAAATGTCAAAGATACAGACAAAAATAATATTGAGGGTGCAAAAGTATCCATTCA<br>AGACATCACAGAAGAAACAACATCGGAAGGATATACAGACGAAAACGGAAATATTCTGTTTT<br>ACGTTCTTGCAAACAACCATATTTCCATAGAAAATGTAACATATGATCTGTATGAAATTAAA<br>TCAGGTCTCGTTTCAATACCGAAAATTACAGAAAATCTAACAGCTAAAATTGTCCTAAAAAA<br>AATAATAATAAAATATAATCTAAGTTTTACAAGTACAACTATGGAAAAAGCCTAATAAAATG<br>ACCCAAGAAAGAGCATACTGCGGGAACTGCATGAAAAAAGTAGAAATCAAAGATGGAAAAAT<br>TACAATCATTAATAAAAAAAAAGTTCTACAAGGCAAATGTCCATTTTGTGACTCCCTCTTAA<br>TAAAAAAAATATAAAAATGGCACTCGAACTTACAAAGGGTGATACCTTAAAAATAAAAATAG<br>GTGTAAAAAACGAATCTGATGGATTTACAGGAACGTACAAAATAAGTGTTGTGATCATGGAT<br>TCTAAGAATACGAAATACTACTTTAGCTACCATACTGACGGAAGCACTTTTTCAACCACTCA<br>ACAATCAATAACCGAAACACTGACGAAAGGCACAGAGTATTCTCTTTCATATTCCGCTGTCA<br>TACCTGATTACGAGGATGGCACAGTCACCGCAAATATAGGAATATACGGAGAAGAGGGGATA<br>ACAGAAATCATCGGAACAGGATGGTTTGACGCATATACTTATAGCAAAAAAAAAATGAATTT<br>GAGCTCGCTATCGGTGACCCTCGAGAAAGGCTAAC |

**Table S3B: Sequence of the non-matching *Metallosphaera* sp. virus probe.** The region colored in grey refers to the polynucleotide used.

| | |
|---|---|
| *Metallosphaera* virus (unclassified archaeal virus, 9.780 bp linear DNA), NCBI Accession no. MF443783.1<br><br>1 Polynucleotide probe (300 bp) from IDT (Leuven, Belgium):<br><br>*Metallosphaera* vir_nonprobe | CCCCCCCCCCCCATGCTGAAAGGGAGGGCGAGACCTAGCCTACCCCCCCCAGAGCAGGGG<br>GTTTAAAAGCCACGTCAGTTTGACGGATCACCACGCAAGTTGAACGCGGGGAGGGTTTAA<br>ATACCCCCCCTAGCATGAAAGCTTATTAACCCCAGGGGTTTTTGTCCTAATGCGAATGAG<br>CAAATTACAGCCCATAATACGCCCTGCCCCGCCCTGCAAGTACACTTTACGACACAAACC<br>GAAAGTAAACCTCAGAGTGAAGGGTTCTCTCCAACAGGGCAAAAAGCATTTTAACCCCCC<br>CATCACGACCTTCCGCAAAATAATTGGTCAGTCGTGGCTAAATTAAGTGAAGACTTTGTT<br>TAGGTAGTTTCGGTGGAGGGTGTGGCTAATAAGGGACAGTAAAGGTCACCATGTCGTCAA<br>ACCAGTCTAACCCAAAAAAGGAGGTCTCCCTGGAGGAGTTTTTTGAGGCCGTGAAGGACG<br>AGGTGATCCCCCTGAGAATGATAGCGGGGGATTACGTAGTCCTAGAGGGCGTCCACATGG<br>AGAACCTCAGGGACGGAAAGAACGGCAAGATAGTTGATGTGGTGGGCAGGATCGTCGTCA<br>CGAACATGGGCCTACCTGAGGGCCAGAAAGTCAGGTTCACGTTAGGCCTCTCGAACGCCA<br>TGGAAGTCGCCCATCATATAAAGGACGGCATAAAGTACACCATGATCAAGAAGGGGGAGA<br>TGAGGATAGTTGTTAATGGGTTTAGCGAGATCCCCAAGAGACTCTAGAGACGCTGAGGTA<br>TACCTAGTCAGGCCCCTGAAGGGCGTTTTTTCCCTCCAAGACCACGCTTTTCTCGTCACG<br>GTACGTGGCGTCTTCCCACGGAAGGGCGAGTTCACTCGAAAGAGCCACTGGGGCCTCATG<br>GTGGACGTGGACGTGGACGGCCTGTCCCTAAGGTTTAGGTCTAAACTCTACTTCACGTGG<br>CAAGTGGAGATGGGCGGTAGGCCGTGGGCTTTCGTGCCCGTGTACGGCTGTGCCGATATA<br>ATCCAGGAGTACGGATACCAGAGAGGTGTGGCCATATGTCAAAAATTGAAGTTGAGCAGG<br>TAGTCAAGTACATATTGGACCATCCTAGTACCTTCCTATGGTACAGGTGGTACCTGAGAC<br>AGAGGCACCACTTACCGTTCTTCTCGCTCACGTACAGTGAGGACTTACGACAGAAGGGGG<br>TGCCAGTGGACATGACCAGGGATAAGGAGGTAGTCGTGTTGGTGCCTATATACACCCCAT<br>ACGGCGTAGGCGGTGCTGAGGTCTTGTTGTCTACCCCAGATAAGGTTATGTGGAGTGACG<br>ACGACTTCGCCTTCTTCGGCAACTGGAAGTTTTACGGGAAGTTCGCGTGGATGGTCTCGC<br>TGAACCGAAAACACCCCCTGAGGCGGTACGCCCTGAACGAGATGAGGAAGGAGGCGAGGA<br>TGTATGCAGGCAAGTCCTGACATGGGGGCATTAACCCGCGCAGGGTATAGAGTGACGGCC<br>CGTTGGACGTGGAACGACGTCACTAGGGCCATGTATTGTTTTGAGACATCTAGGACAGCT<br>AACGACTTCGTGGGGTGCGTCAAAAAGCGTAGGACTGACGCCCTCGCTGTCTACCGATAC<br>CTACGAGACGTCCAGTCCCATCAGGGACTGATATATGAGGTACTCCCGCCCGCATTCCAG<br>GGCAACCCAGAGGAAGAGGAGGGGGAGGGGGAAGAGACTTGTATCACCTTCGTGGTCTAC<br>GACACGGGGTACTTCTTCCTCTGGGTAGACCCTAACCAGATCCTTCAAGACGCCTTAGAC<br>GCTAACGCGACCATGAGGATCAAGTACCCTGGGGCCTACCGTGTCATCACCACGCCGTAT<br>GACCTAGATTTCTCTCAAATAGCGTGGAGGCCCATAGTGCCGTCGGGCGAGGACTACGTG<br>ATATGTAGTAACATGGACCGTGTGGACTTCAGTGACGCGATAGACAGGTACGTCAGACAG<br>CTGAAGGCGATAATGGGGGCGATATCATGATCACGTTAGACCACGATGTACCATATGACG<br>GGTATAGACATTCGGCGAAAGGGAAACTCGTCGTTGAGTTTTGCAAGTCGTTCGAACTGA<br>GGTGTTGGTATAGGCGTTCGTCTACTGGGAACGTCCACGTGGCTATCGACGTGGACGTGG<br>ACTTCCTCAGAGAGTTGGAAATCAGGGCCGTCTTACTCGACGACCCCATGAGGATATTGA |

| | |
|---|---|
| | ACGACTTGCGTCGGCACGCATTCCACATGCCTACTGGACGGCTCTGGGACGTGAAGGTGG<br>ATAGGGAAGGCAAAAGGCAGGCGGGCCAGTGGGAGGTCTTGTTCTCCCCTGAGTAGGGTT<br>TTTTACTCCACATATCCTACTTGTATCGTATGAGTCAAAGTCAGCCTACTCAGGAACAAA<br>CCCCCCCACAAGAGGAGGGGAAGGACGAGAGCCAGTGGACGGAATGTGAAGTCTGTGGGA<br>AGAAAGTGCGGATGAACATGTACAAACGTCATATGTCTGCGGTGCATGGGTTCATTACGA<br>AAGAGGAGTACCAGAGGCTCAAAGAGGAGGTGGAGAAGACTATACAATCTACGATACACA<br>GTGCACTCCAGGAGCACTGGAAGGGGCACTACAAAAGCTTAAGCGAACTTCTTGAGGAAG<br>GGGAAAAACATGCCGAGACGTGTCCTACATGCCAAAAGGAGTTGGAGGAGTGGCTCAAGA<br>AAAAACGGAACGAGAAATACAAACCCGCTAATAGGAAGTGGCTCTAAATGACCAAGAAGA<br>AGAAGGAGGAACAGGAGGCCCCTAAGGAGGAGGGGCAGGCCACTCCCGAAACCCCAAGTC<br>CTGAAACCCCTCAGGAGCCTCAGAAGGAAGACCCCGTTCAGGTCATCATGAGGCTGACGA<br>CCGTAATACAGGAACAGCAGAAACGTATAGACCAGTTGGAAGCAAACCTCAACAGCGTGA<br>TGGAATTCCTGAAGGCACAATCCCAGGCACCCCAAGGTCAGGGAGGAGGAGGGGGTCTAG<br>TGGAGGCACTAGTACCCCTCTTAGGGAAGTTGATGGAGCCTTCGAAAGACCCCCTACGGG<br>ATATCGCCCTAGAACTAATGATCGACAGCATGAAATCGAACGTTGAGCTCTCGAAGGCCA<br>TAACACAGAAGATCATCCAAGGGGTGCAAGACCGAGTCGCGAGGAACGTCGTGGAGACTG<br>TCGCCACGGGGGTGATTACACACGAATGAGCGTATTAGAATTATCAACTTAACAAGGAAG<br>TTAATGATTGAACTGGGGGTGGGTACGTACTACCCCATGGTACAATACCGTCTTACCAGT<br>CTCCCAGAGGAACAGATAAAGAGGGCGTATAAAATCATAAGGGATGAGGTGAAAACGTGG<br>CCATAGTCCATGAGGAGGTAACGTTTCATACTGAAGTGGATGATGAGAGCGTAGAGGCAA<br>TCAGGAGGATCATAAGGGACCAGTGTGGAGAAAACCCAAAGCCTGAATGTTGGGGCCCGT<br>TGGTGTGGTCCATGTTCGACAGCGTGGCCCGTGCCATCCCGTGCCCAAAGTGCCGAGGGG<br>AGGCACTGGAATTCCTGACCTTTTTGCATGATTTAATTAACATACAGAAGGGTAGTGCTA<br>TATACAATACCCCGCAATTCCTTAGGAAGAAAAAGGAAGAGATACTGAGCCTACTCAGGA<br>CCCGAGGGATATAATATAAATAGGAGAACTGATGTAACTATGGTAGAGAGCCATGGTGTC<br>CAATAGAGAACTCGCCAAAAGGCTCAGCCAGTCGGCCAGGGAGGCGTGGAATTCAGAGGA<br>GGGCATGAGGATCAGAGAGGAGCTAAGTGCAGACTCTGCCTCCTATGGGGCGTGTCTGAG<br>GAGAGTCATGGAAAGTGGAGGAGGCAGTCGTGGTTACAAGGATTGCGCGAGGAGGTCAGG<br>AATAGGTACGGCGTTTGCCAGTGTCTGGGGGTCTACAAAGGCCTATGGTAGGCCGATGGT<br>GGAAGTAGTAGCGAGGTGACCTAGGTGGACTTCTCCAGACAAACGTGGGAGGGGCCAGTG<br>GCTGTTGGCGTCGGCGGGTACGTGTGGAACAAGGTAAAGGGGAACACGCAAGTTCAGACG<br>TTCGTGAGCAAGGTAGGTGGCAAGACGAATGCAGGGCTCATCTTAGTGGCCCTAGGCGTA<br>GTCGCTAAGGCATACAAACCCGAGTCAGAAGCCATGAATATACTGGGGTACCTGTTGGCG<br>GGACTCGGTGCGGGGGCACTCGGTGACGACTTGCCCGCCACTGGGTCACAGGCGACGTTT<br>ACGAATAAAAGCCCAGAGTACCCTAGTCCAGTTAGTGGGGTGAACGTGCTGTGAAACTCT<br>ACATGAAAACGATTGAGATCCAATATTCCATCCCTGCGAGTGCACCATCCGAGTCACAGG<br>TGCCTGGAGTGTTCATAGACCTCTCCAGTGGCACTAGCCAAGGCCAACTCCAAGTCCCCG<br>TGAACCAAGAGTGGGTCATCATAGACGTCTACAACAGGGGGTCTCAGGACGTCGGCGTAG<br>ATGCCGTGGCGACGTTCACTAAGAACAGCGTCTCCAACGTCCTCACTACGGACCCAGTCT<br>CATCCCTGAACATCTCGAACCCATCCAGACCAACCTACCCGCCAATACAGATGGCCCCAG<br>GGACGATATTGACCTCCAAAGTCACTACGCTCACGTCCAACGGCACTAGTGCGGCCGTGG<br>ATAACCTATTCATAAAGGTCAAGATCATAGACTACTCGCAGTAAACACGCGTAAGACATC<br>TCCTTTTTTAGTAGTATTTCCTTCTTTATAGTATGGCATGCCAGACTATACAAGACCTAT<br>CGATAGGCCCGTACACCATAGAATCGGGTCAGGTCTGTGTGGAGGGCACTACCTCAGGGA<br>CGGCTAGCCTGACTATGAAAGTGGGGCCGTTCACGTACACCAAGACTATCCCATGGGGGG<br>CGGTAAACCCCCAACAGACGTTGGGTACGTCCTCAGGGAGGGTCTACCTTATCGGGACGA<br>ACCCCCTGTCGTCACTCCTAGTGGACGACGTCCTAACGGTGGCTGAGTATGCACAGAAAG<br>TGGCGTCGTACTTAGGGTACAGCGGTTCGTTTTCCATAGTGAACGTGGGTTTGACGAATG<br>GCTCAGACTGCAACGACTGTGTAGTTTACGTGGACTACGTCGCCTCTTCTCCTCCTCTTG<br>CGGTAGCCCTAGTGATCGCTCTCGTGATCGTTTTCGTCATAGCTATCGTATTGGGTATCT<br>ATTTCATTTCGGCCGCGATAGAGAGTTTTCAGCCCGCACCTCCCACGCCCCCTCCTCCAG<br>GGAGTCCCCCGTCCCTGTACCAACAGTACTACCAGGAGTACGCCCAGTATTTGAAGGAGA<br>AACAAACGGCCTCAATCACGGGAGGGGTATTCGGCACGTCCACGGCCCTAGTGGCGTTAG<br>GGATAGCCATTTTAGCCTTCTTGGCGTTTACTGACCATGGTGGGAAGTGATGGTGTACCA<br>GATATACGACGGGCAATACACGGTTTACCCCGTGGGAAACCCCCAGGTCACGCTCATCAG<br>GGTAGACGTAGCGGGGAGAGTGAAAATCATGATCCTCCCCACGGTGTCAGAGGCCCTTCA<br>ACTGACCCTGAACGGAAACTCAGGTTACCTGAACCAGGGAAACGTCCTCTCGGCGGGAAA<br>TTGGTACGAGTTCGAATACGTGAGTTGTGGGAAAGACATCATAAGCATAAATGGACACA<br>GCAAGGAGTTTACCTTAGGGTGATAGTGTATTGATTAGAGGTCAGATACACAAACTCTCC<br>AAGTGGGAGTTAAGGGACTACGCTAGGACACTAGTCCCCGTCTTTGAATCCCACGAGGTG<br>GACGGCACTAGGGACATGAAAATCGTGGACGCCGTGATATCGTACGCCATGAAAGGGACG<br>AAACCCTGGCTGAGGAGGGGGTCAGCCCTCCAGGGCTTTATAGGGCCTAGGATGTCTGTC<br>CTCGAGGGTTACCTCATCTTCCCCACGTCAGGTATTGAGCCCGTGACCATCCAGTCCACT<br>ACCCTAGAGAACTCCTACCTAGTCCTAGATATCGTGGGGGCAGGTGGAGGAGGTGGTGGT |

| | |
|---|---|
| | GGAGGTGGTGGGAACACCACGGCCACTACCCCGTTCGCCCAGGGTGGTGGAGGTGGGGGA<br>GACGGTGGCCGTCTCATTATCAGGACCCAGGGGCCACTAGTCTCTGGGACCCAGTTCACA<br>GCCCAACTGGGAGGGCCAGGTGGGACAGGTGGAGGGACAGGTGCCACAGGGACACCAGGT<br>TCAGCACCAAACGGGGCGACTCAGGTAGTCCTAGGGAACGGCACACTCACGCTCAGTGTG<br>TCAGGTGCGTCAGCCCCTACCATGCCCCCGACAGCAGGAGGACAGGAGGCAGGAGGGTCA<br>GGAGGTGCAGGAGGGTCAGCAGTGTTCACGTACTTGCCAACGACGCCCATATTCTCGGCC<br>ATACAGTTGAACCCAAGGACGTCCTTCCTCTCGAATTACTCGGGCGTACTGGACCAGGGT<br>ATTGGTGGAGACGGCGTTGGGTCTGAGTACTTCACCCTGTCCAACGCCACGCCTATATTC<br>GTCTCGACGCCACAAGGAACGAGGGTCACTGGGGGAAGCCCAGGGGGCACAGGCACCTCA<br>GCCACGGCGAACGTCCCCTCTGGGGCAATCCCGCTCAGGGGGTCAGGTGCAGGAGGAGGA<br>GGGGGACAGAACTCCTCTTCCGCCCCAACAGCGGGAGGGAACGGCGGGTCTGGAGGGCCT<br>GGACCTATCATTATCTTGGTGATAGGATGACAGACTTGGTCACCGTGGCAGTGTTAGGGG<br>TACAGACGGCCGTTTTCATCTACGCTAGTGCCAAACTGAGGAAGATGAGCGAGATGGCCG<br>ACAAGTTGGCGTCAGCCCTGTCCAACATAGCAAAGGAGATAGAGACGGCGTCCATGGACA<br>TGAAAAGCATACAAAGGGAGATGACAGAACTGGAAGACTTAATTCGGACAGCACTGAGGG<br>AATAGGTATGAGCTGTCAGACATGTGGTCAGCAGGGCGTGTGTTACGTCCTGGAACACGT<br>GGGTACTGAGCCCATCTATATAAACGGGCAACAGTACGGCCAGGGCGACTTTTGTTTCTC<br>CTCCCCTGAGCTCACGATAGAATACCAGGGCGTACAGTACCAGATCGACATGTCCTACGC<br>CCCCACAGTGCAGAACATATACGGGTGTTGTCCCTCAGGTACCACAGCCAAACTCACGAC<br>GGTGACTACGATACAGGGACAGAACACTCCCATATTCGTCCCGTATGCCGTCCTAGTGTC<br>TCAAGGTACCACGTCCACTAGTAGTACCACGCCTAGTGTCCTAAGTAAGGTACCGTGGTG<br>GCTCATCATACTACTCGTCATACTGGGGGCTATACTCCTAGGAGGAGGAAACCCATGAAT<br>TACACTTGGTACGTGCAGAACCAAGACCCCTCGACGATCCAGTTCTGTGTCATGCAGTCT<br>GGAGGGTATAACAACTGCACCTCAATACCAGGGAACAGCACAGAGCCGTTCACGATACCA<br>GACATGGACGTGCAGATATATGGCACCCCCACGGGGTACTACGTGTACTACTCTAGCGTT<br>GGGGTGAAGACTAGTACCAAGTCCCTCCCGTCATCAGGATCAACGATCACGGTGCCCCCA<br>TACACGGGATGTGACCCCCTACTCACGCAAGGACAGGGGTACCCCCCACTCGTGCCGTGT<br>TCGTCCCCGTTGATCTTCCTGAAGAGCGAGAGCACCCCTGCCATGTTTATCGAGAACTGG<br>GGGGCAGAGACGTCAGGGTTTTCATACGGGTACGGGATAGCGGGGAGCCAATTGGCCATA<br>GCGACAGGGCCAAACTGGGTATACTACCGACAGTACCCCCAGGCACAGGTATACCTTATC<br>GGAAACTGGAAGGTGGTGAACGGAAACTACGTAGACATGGACTCGGGGAAAACGTTCTGT<br>GGGTGCCAGGGGAAGAGCGTGTACGTCATACCCACGTCTGACCTACCTGGGCCGTCCTCA<br>GGGACTACGTCCACAGGGACTAGTACCACGGGGACTTCCACGAGTACAACAGGGACTCCC<br>ACAGGGACGTCCACTCAGCAGACGACACAGCCCACGACAGGTATCCTGAACGTGGCTAAC<br>CAGTGTTCGACGGGGGTAGTCGTGTACGCGGGAGGGAAAACGGCATACGCACCCCCATAC<br>CATTCCATCAGCCTAGAAGTACCCATCCCCACGGGGTATACGCTTATTAGCCAGGGTCAG<br>ACAGTTGGTAAGGGGACGGTGTACCAGAGCGGGGAGTACGTGTCCCCAAGTTCTTGTCCC<br>TCTGTTAGCCCTACACCCAGCCCTACCCCGACACCGTCCCCAACTGGGGGACAGACCACT<br>TCCACGACTACGACTACCACGACTACCCAGACCACTTCAACACAGACTACCCAGAACACA<br>CAGCCCCCAACGACGCCTAGCATGTCGTGGGGTCTTATCCTACTCTTGGTAATCCTGGGG<br>GCACTAGCCCTCTCAAGTTAAGCTTTTTTAGTCCAGGTCTCCCATTATACCTGAGCGGTG<br>CGAGGTGGATGTGACGGGAGGCTCTGGGAATGTCCGAGAGACGGAGGCGTACCCGTCCTC<br>CACTGCACCGCACTAGGTGAGTCACATGAGCCAAGTCCCAAAGAAGAGACTCGGGGCAAG<br>GGTCATCAGAGCGGTTTTCAACGGGGTCTCCTTAGGCATCCTAGGAGGGGCAGGAGTGTA<br>CGTCCTAGCCCAGGGGGTGAACACGTTATCAGGCACCACTACTTTGAACCCTACGGCGTT<br>TCTCCTGTTGGTCTTGGGGGCGTCCATCACGGCCAGTGTGGGGATAGAACTCTCCAAGGA<br>CCTAAGCGGTGAGTAGTCTCAGTCAAGTCTCCTTTATTTTACCCCCTGAACCAATAGACC<br>CAAAATGGATGTCCAAACCCAACTCAAAGAATGGATTTCCCTGTTGGACGACAGGCTGTC<br>GTCCATAGAAATGAGCCTCAAGTCCCTCCAAAAGAGACTTGAGCAGGCAGAAAAGCGACA<br>AGAAGAACTTGAGGAGGCGTTCGACCAGCTGAAAGAAGAAATAGACGACTGTGTGGCTGA<br>CACGACCTACCTCCACGACGTACTGGAGGAGGTCCAAGACGACATAGACGACCTAAGGAA<br>GTGAGCCCCGTGGAGTTCGTATACAAGATCAACGACGACACGGTCACAGTGTGGGTCACT<br>GACGAGAAGACGGGGAAGAAAGTGGCGTCCCTACAAATGCCGAAGAAGGACGTGGACTAC<br>GTCGCGAGACGTGCAATACAAGAACTCGACATAGCCCCAGATGAGGCATACGACTTAGCG<br>GTCTACTTCACCTACGTTAAGGTCACCTGGGGCCTCCTCTGAACCTCTTTTTTCTCTCGT<br>ACTTACCCCTGACCCTACGGTCAGGCTCTACTTCTCTCACGAACCTGACTCCCCCATCCT<br>CATCAAACATATGATTTATTTCACAGCAGGGCTTTATATAAATTGCGGAAGGTCGTGATG<br>GGGGGGTTAAAATGCTTTTTGCCCTGTTGGAGAGAACCCTTCACTCTGAGGTTTACTTTC<br>GGTTTGTGTCGTTAAGTGTACTTGCAGGGCGGGGCAGGGCGTATTATGGGCTGTAATTTG<br>CTCATTCGCATTAGGACAAAAACCCCTGGGGTTAATAAGCTTTCATGCTAGGGGGGGTAT<br>TTAAACCCTCCCCGCGTTCAACTTGCGTGGTGATCCGTCAAACTGACGTGGCTTTTAAAC<br>CCCCTGCTCTGGGGGGGGTAGGCTAGGTCTCGCCCTCCCTTTCGGCATGGGGGGGGGGGG |

**List of Supplementary Tables that are provided as .xlsx:**

**Table S4: CRISPR Cas systems of highly abundant Altiarchaeota.** CRISPR-associated proteins based on UniRef100 [52] annotations in assemblies and based on searches for a combination of the key words “Alti”, “Cas”, and “Alti”, “CRISPR”. The table also contains information on CRISPR arrays and associated Cas proteins detected by CRISPRcasFinder [54] in Altiarchaeota bins. ND=non detected.

**Table S5: Virus Orthologous Groups (VOG) on Altiarchaeota virus genomes.**

**Table S6: Variants derived by mapping reads from MSI biofilm (BF) 2012 and 2018 samples to 2012 genomes of Altivir_1_MSI and Altivir_2_MSI, respectively.** Details on single nucleotide and other polymorphisms are given together with the information if variants affect a region matched by a spacer. Analysis has been conducted in Geneious Prime version 11.1.5 [101].

**Table S7: Protein annotations of open reading frames on Altiarchaeota virus genomes**. Annotations were performed using HHpred [61, 62] and UniRef [52]. For the most abundant viruses, e.g., Altivir_1_MSI and Altivir_8_HURL, PHMMER (PH) [64] and DELTA-BLAST (DB) against NR [63] were additionally used. MSI=Mühlbacher Schwefelquelle, Isling, ACLF=Alpena County Library Fountain, HURL=Horonobe Underground Research Laboratory, GA=Geyser Andernach

**Figure S1: Classification scheme for viral scaffolds**. A) Bioinformatic tools (VirSorter [82], VirFinder [102], VogDB [103] (version Vog93, e-value cut-off $10^{-5}$), CircMG [104] and End Matcher (https://github.com/ProbstLab/viromics/tree/master/Endmatcher) used for detecting viral scaffolds; B) Combination of results and how it leads to classification of predicted viruses into putative viruses and viruses. Modified from Eßer [105]. VirFinder [102] hits were false discovery rate-corrected using the package q value [106] in the R programming environment [74] applying a p-value cut-off of <0.05.

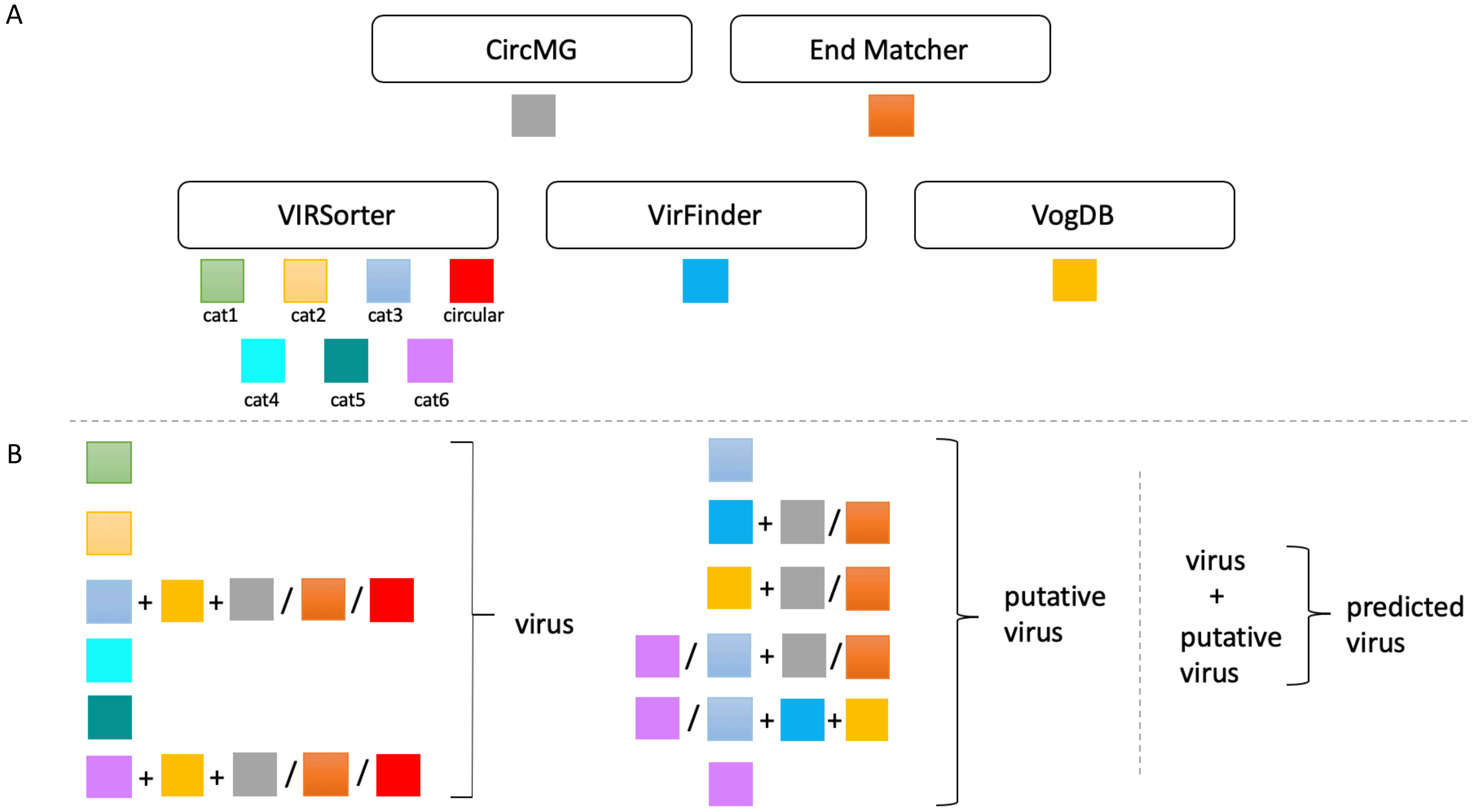

**Figure S2: Rank abundance curves for metagenomes based on ribosomal protein S3 indicating dominance of Altiarchaeota in the MSI ecosystem**. for A) MSI_BF_2018, B) MSI_<0.1µm_2018, C) MSI_BF_2012, and D) MSI_>0.1µm_2018 (stretched over two panels). Assemblies for 2018 samples were performed with MetaSPADES 3.10 [50], whereas for the 2012 assembly MetaSPADES 3.11 has been used.

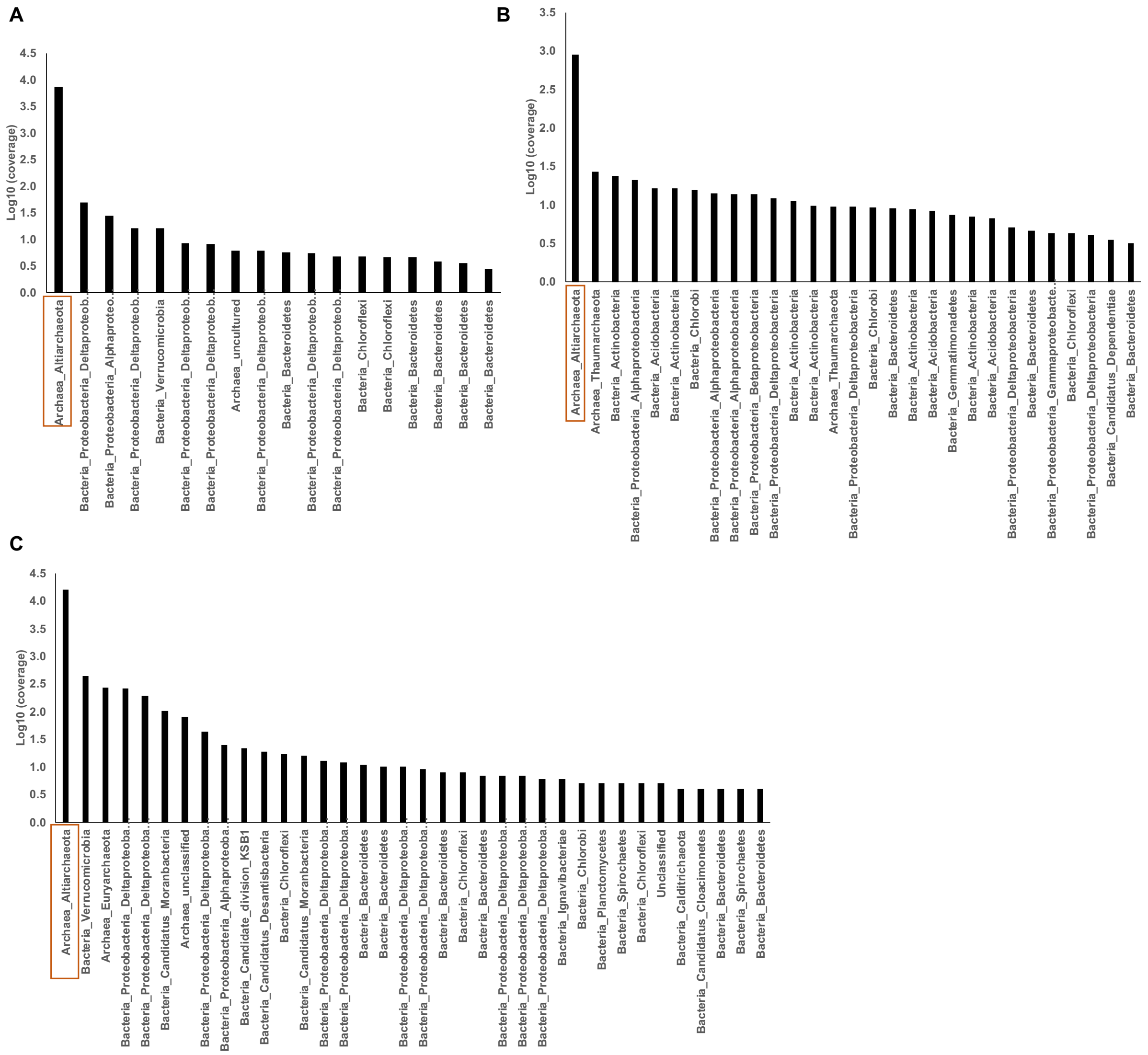

D

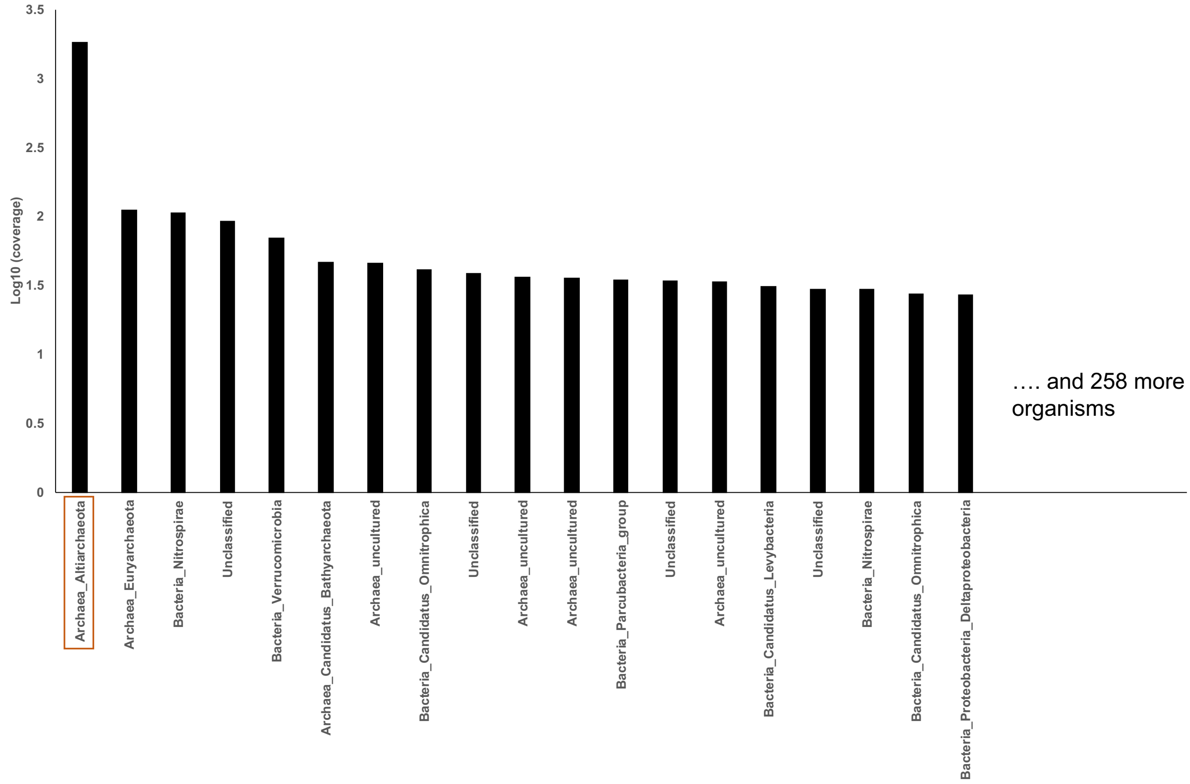
3.5
3
2.5
2
1.5
1
0.5
0
Log10 (coverage)
Archaea_Altiarchaeota
Archaea_Euryarchaeota
Bacteria_Nitrospirae
Unclassified
Bacteria_Verrucomicrobia
Archaea_Candidatus_Bathyarchaeota
Archaea_uncultured
Bacteria_Candidatus_Omnitrophica
Unclassified
Archaea_uncultured
Archaea_uncultured
Bacteria_Parcubacteria_group
Unclassified
Archaea_uncultured
Bacteria_Candidatus_Levybacteria
Unclassified
Bacteria_Nitrospirae
Bacteria_Candidatus_Omnitrophica
Bacteria_Proteobacteria_Deltaproteobacteria
…. and 258 more organisms

**Figure S3: Two distinct direct repeat sequences of CRISPR arrays derived from Altiarchaeota and formation of secondary structures.** A) Alignment of CRISPR repeat sequences from Altiarchaeota genomes obtained from different habitats (Table S4). All direct repeat sequences end with a AAA(N) motif. Alignment was performed using MUSCLE [107]. B) CRISPR RNA (cRNA) secondary structure predictions of the different repeats and their orientations predicted by RNAfold [108].

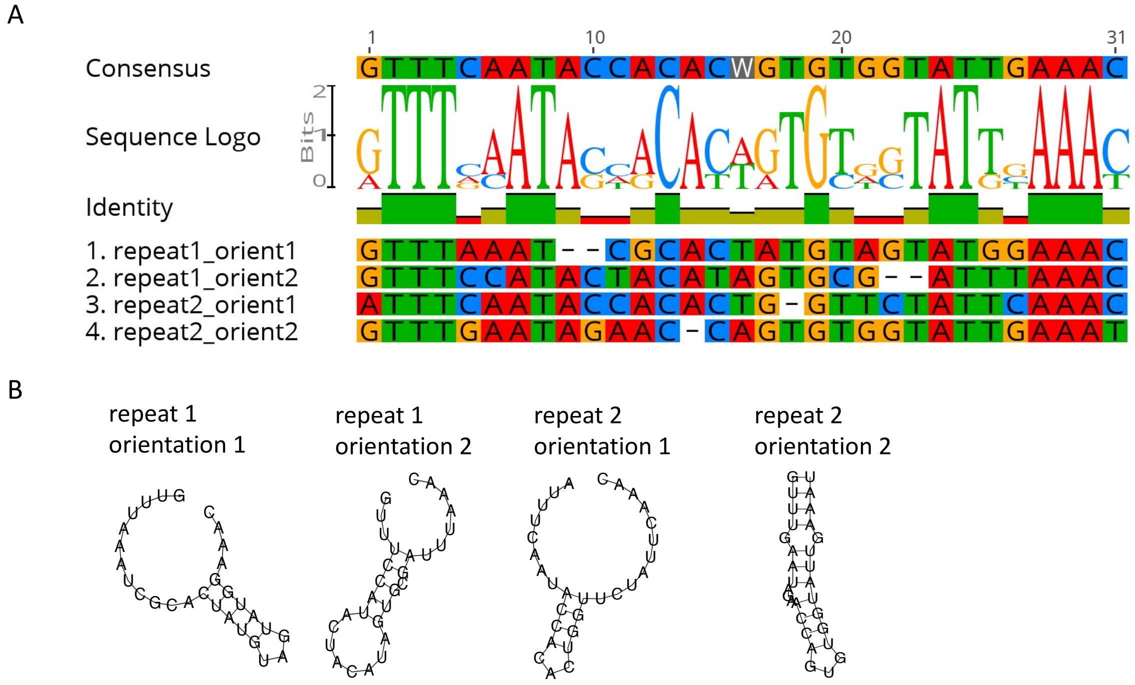

**Figure S4: Viral clusters of Altivir_1, _2, and _6 and their position in the viral network**. Altivir_1_MSI genomes form a cluster with Altivir_6_ACLF while Altivir_2_MSI genomes form a cluster on their own. The network shows how genomes cluster according to similarities with the database "ProkaryoticViralRefSeq94-Merged" [70] after using vConTACT2 [68, 69]. Nodes and edges represent viral genomes and genome similarities, respectively. Visualization was done in Cytoscape [71]. All remaining genomes of *Ca.* Altiarchaeum viruses were excluded as unclustered singletons.

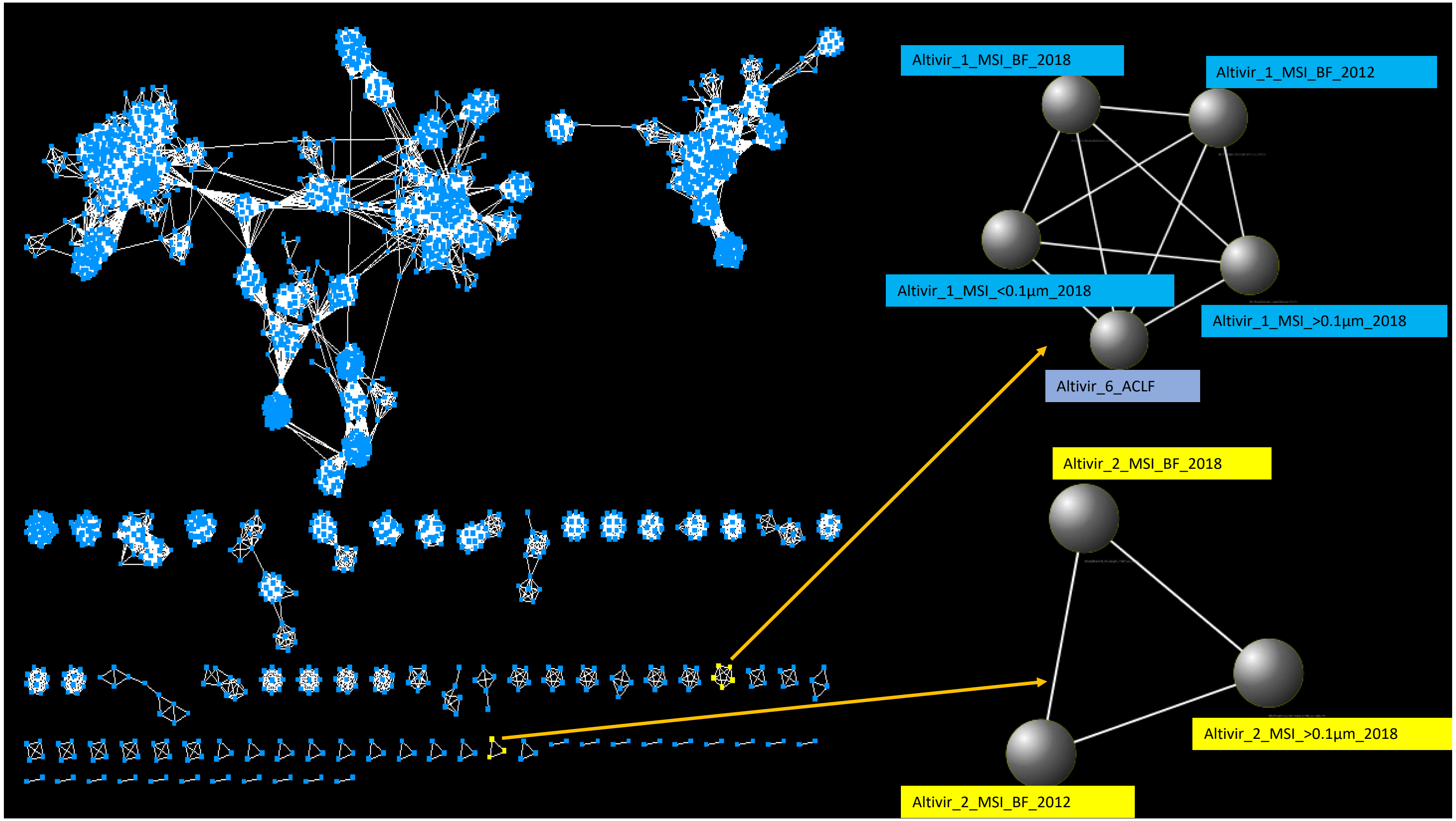

**Figure S5:** Intergenomic similarities between Altivir genomes calculated by VIRIDIC [72].

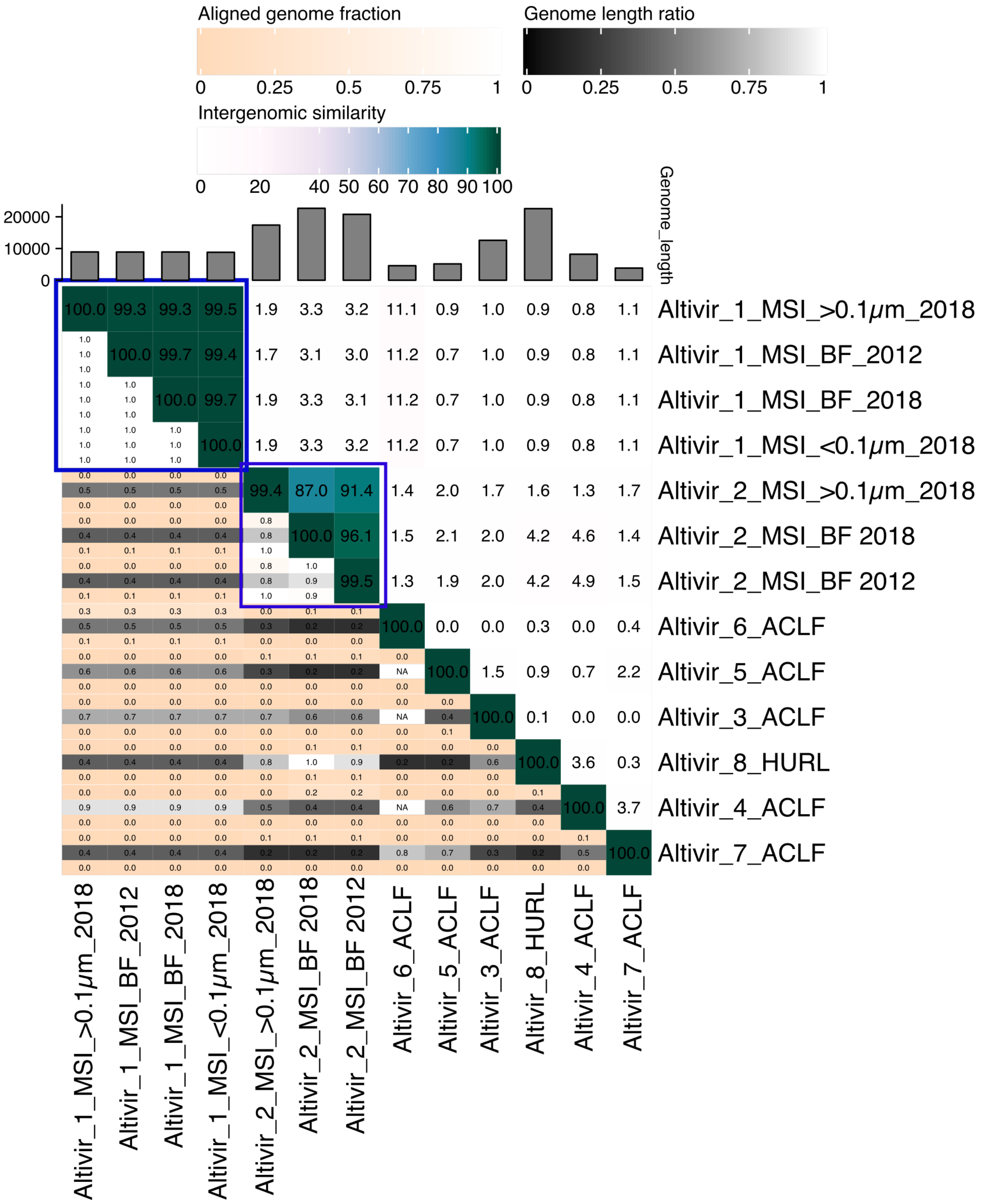

**Figure S6: Development of CDS from 2012 and 2018 samples for Altivir_1_MSI (upper panel) and Altivir_2_MSI (lower panel).** Variant regions can only be shown for Altivir_1_MSI due to the huge gaps in beginning and end of the Altivir_2_MSI alignments. Alignments were performed using MUSCLE [107], and variant calling was performed with default settings of Geneious Prime version 11.1.5 [101]

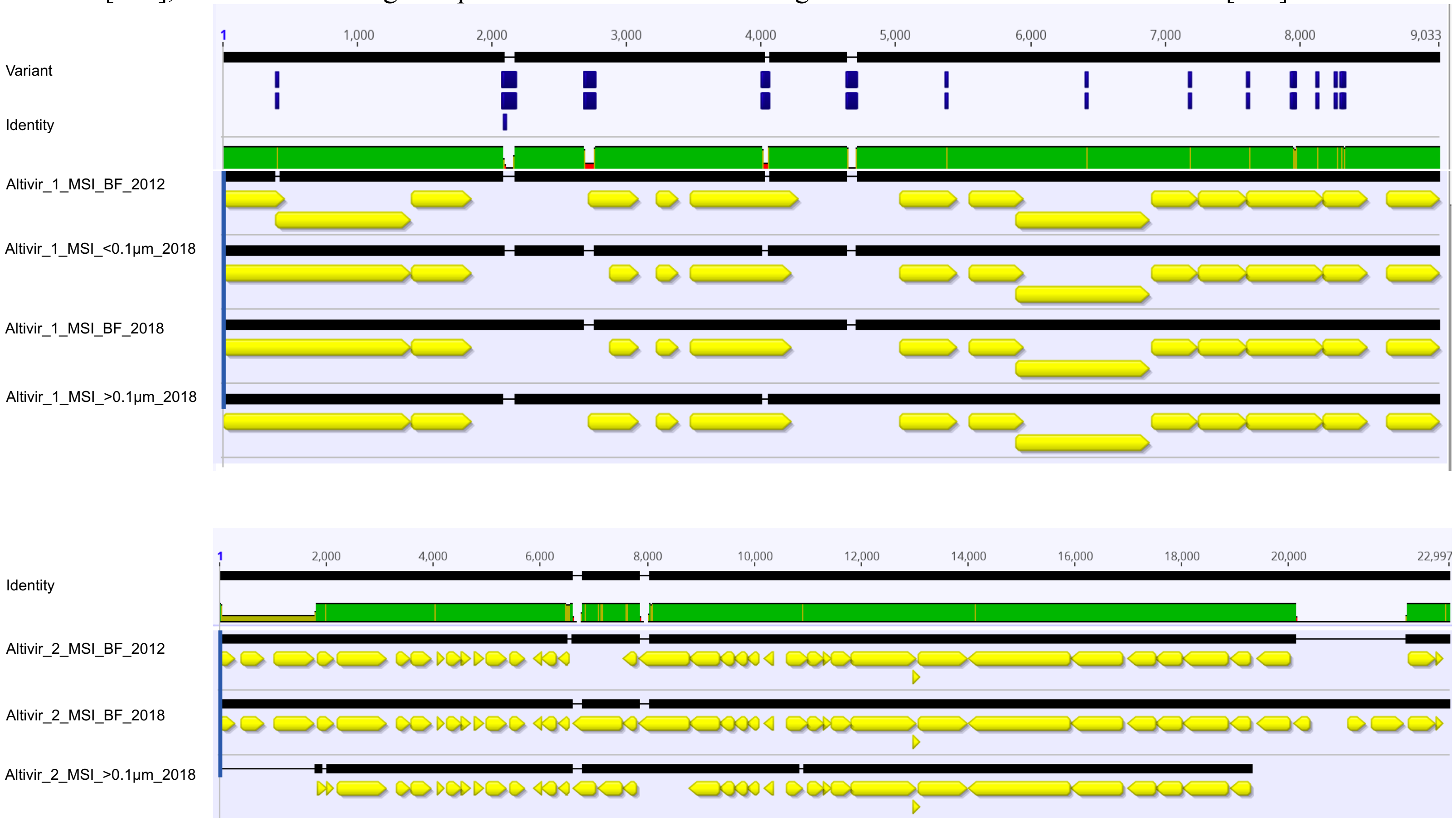

**Figure S7: Spacer mapping to predicted viral genomes and variants therein.** A) Spacers extracted from MSI_BF_2012 metagenomes mapped against genomes of Altivir_1_MSI_BF_2012 and B) Altivir_2_MSI_BF_2012. C) Spacers extracted from MSI_BF_2018 metagenomes mapped against genomes of Altivir_1_MSI_BF_2012 and D) Altivir_2_MSI_BF_2012. Variants (abbreviated to SNP) are based on read mapping from reads of MSI_BF_2012 and MSI_BF_2018 samples (see Table S6) and coding regions (in yellow) are shown. Variant calling was performed in default settings of Geneious Prime version 11.1.5 [101]. Mapping of paired reads to viral genomes was conducted using Bowtie 2 sensitive mode [75].

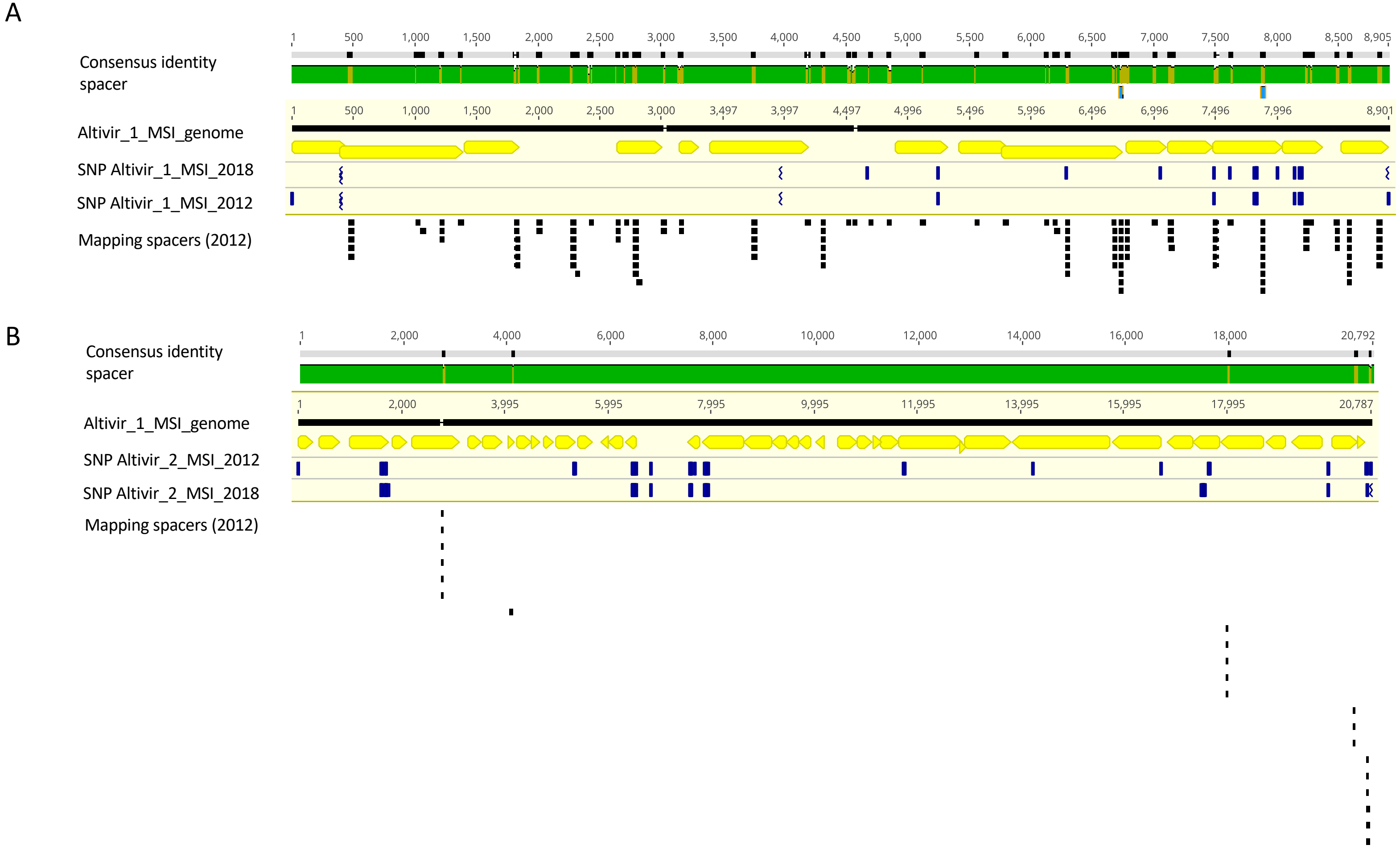

C

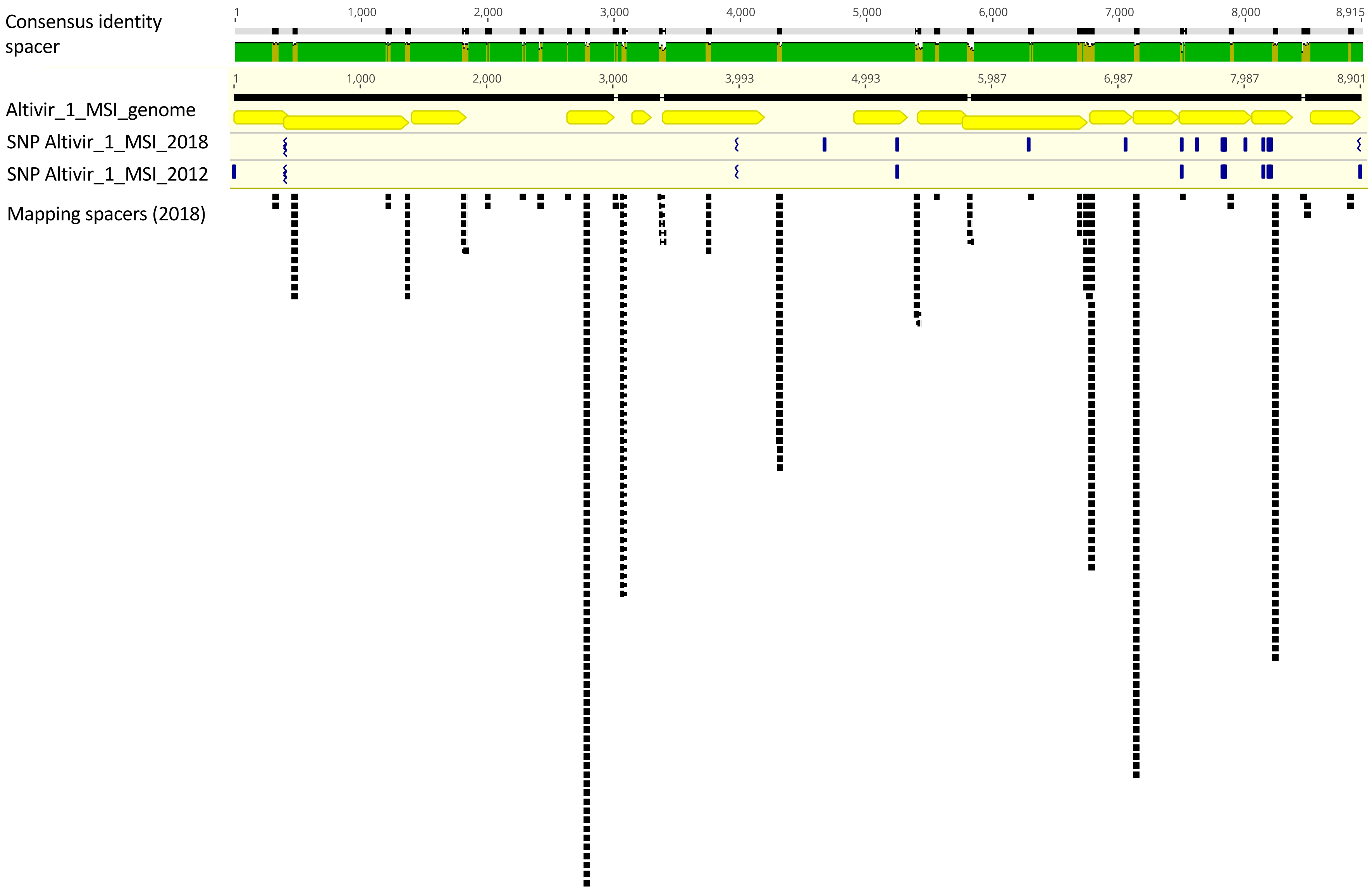
Consensus identity
spacer
1
1,000
2,000
3,000
4,000
5,000
6,000
7,000
8,000
8,915
Altivir_1_MSI_genome
1
1,000
2,000
3,000
3,993
4,993
5,987
6,987
7,987
8,901
SNP Altivir_1_MSI_2018
SNP Altivir_1_MSI_2012
Mapping spacers (2018)

D

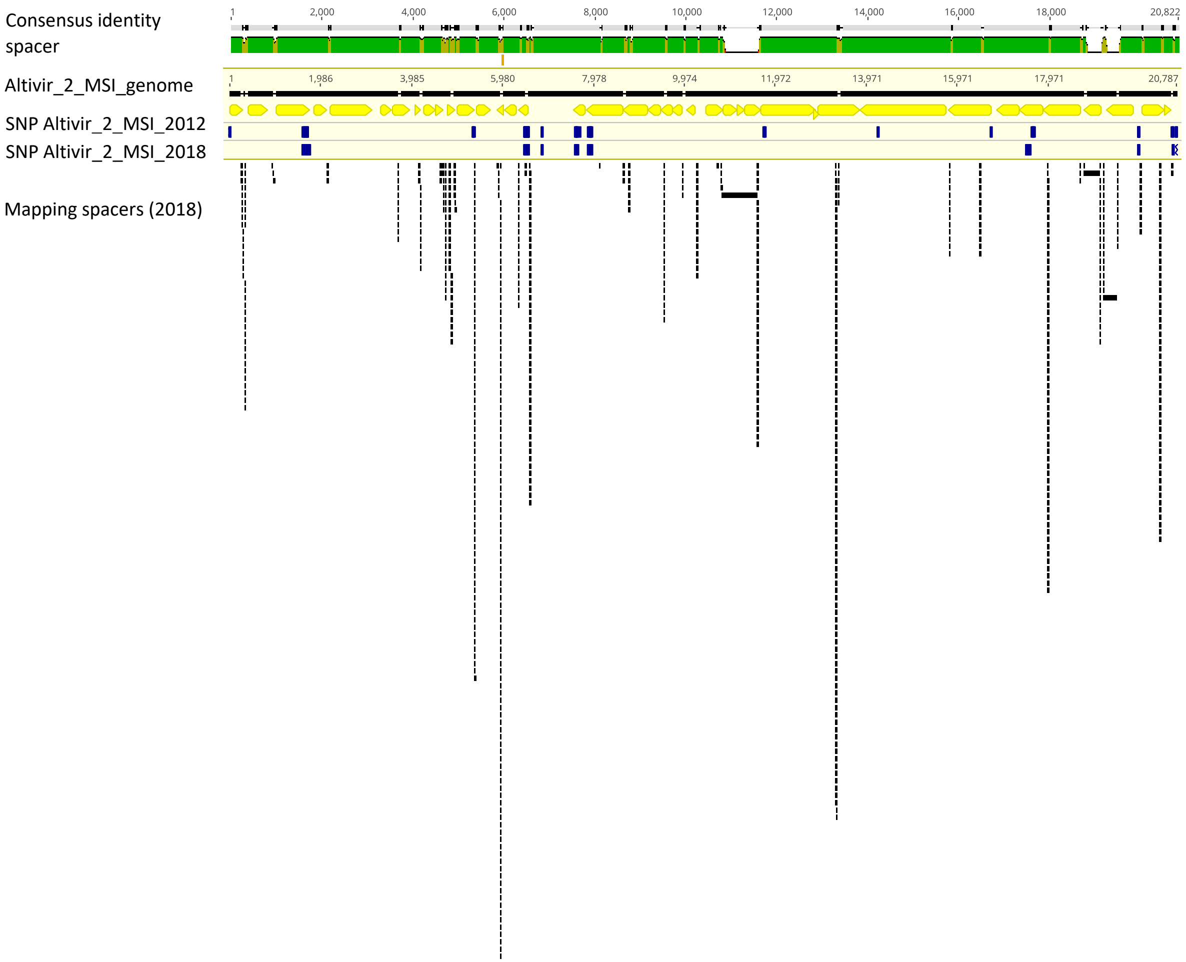

Consensus identity
spacer
Altivir_2_MSI_genome
SNP Altivir_2_MSI_2012
SNP Altivir_2_MSI_2018
Mapping spacers (2018)
1
2,000
4,000
6,000
8,000
10,000
12,000
14,000
16,000
18,000
20,822
1
1,986
3,985
5,980
7,978
9,974
11,972
13,971
15,971
17,971
20,787

**Figure S8: GenomeFISH performed on Altiarchaeota biofilms (BF) for visualization of viral infections caused by Altivir_1_MSI.** For genomeFISH, a chemically synthesized probe mix targeting Altivir_1_MSI was used. BF material was analyzed with A) DAPI (blue, cells), B) ATTO 488 (green, 16S rRNA signal) and C) Alexa 594 (red, viral genomes of Altivir_1_MSI) and D) merged.

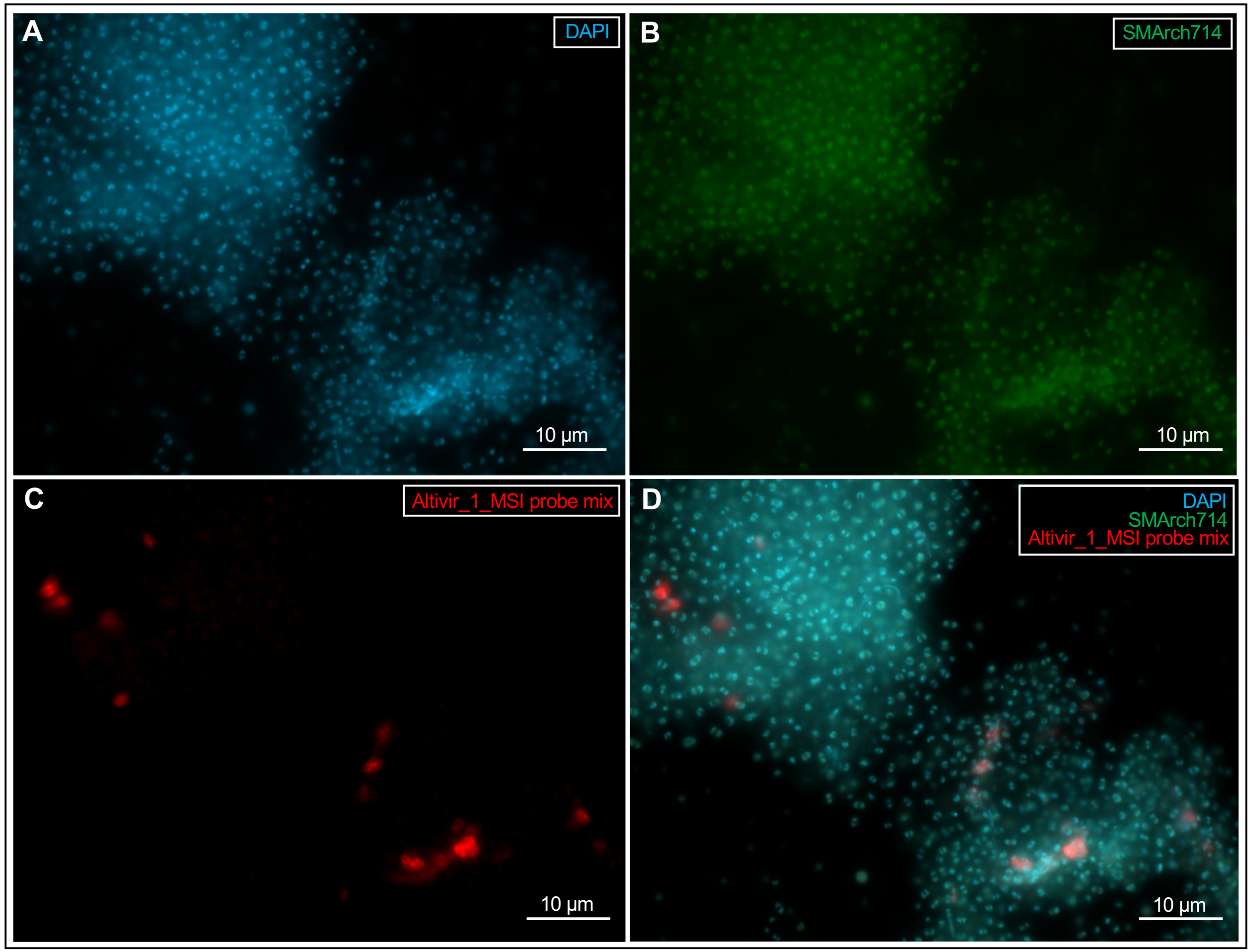

**Figure S9: A chemically synthesized *Metallosphaera* sp. virus probe was used as a non-matching probe for genomeFISH showing no viral infections within an altiarchaeotal biofilm (BF).** BF material was analyzed with A) DAPI (blue, cells), B) ATTO 488 (green, 16S rRNA signal) and C) Alexa 594 (red) and merged (D).

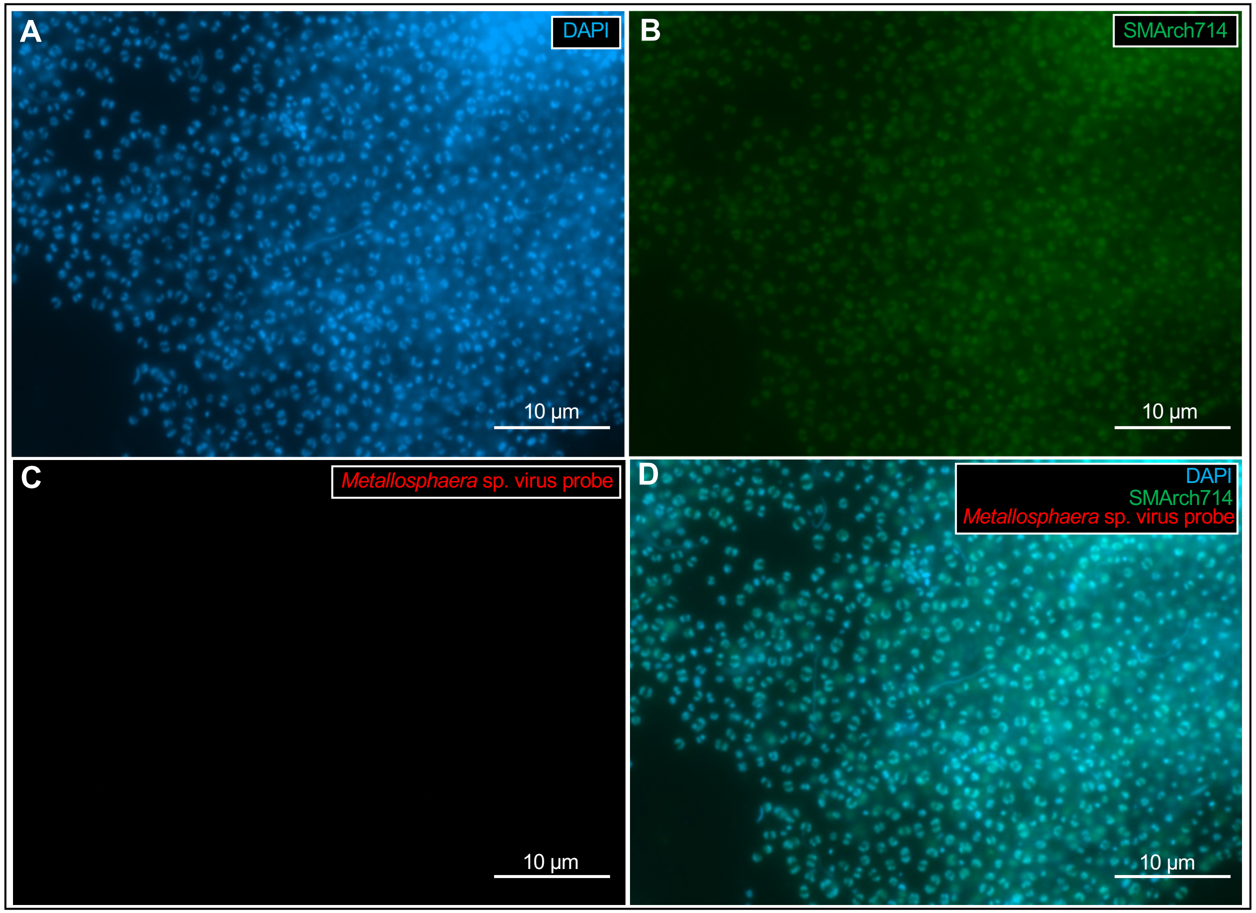

**Figure S10: Bar graph depicting the number of spacers found as singletons compared to the total number of different spacers per sample from the MSI site**. An increase by ~20% from 2012 to 2018 is identified, which indicates a diversification of the CRISPR array in Altiarchaeota and of their respective strains.

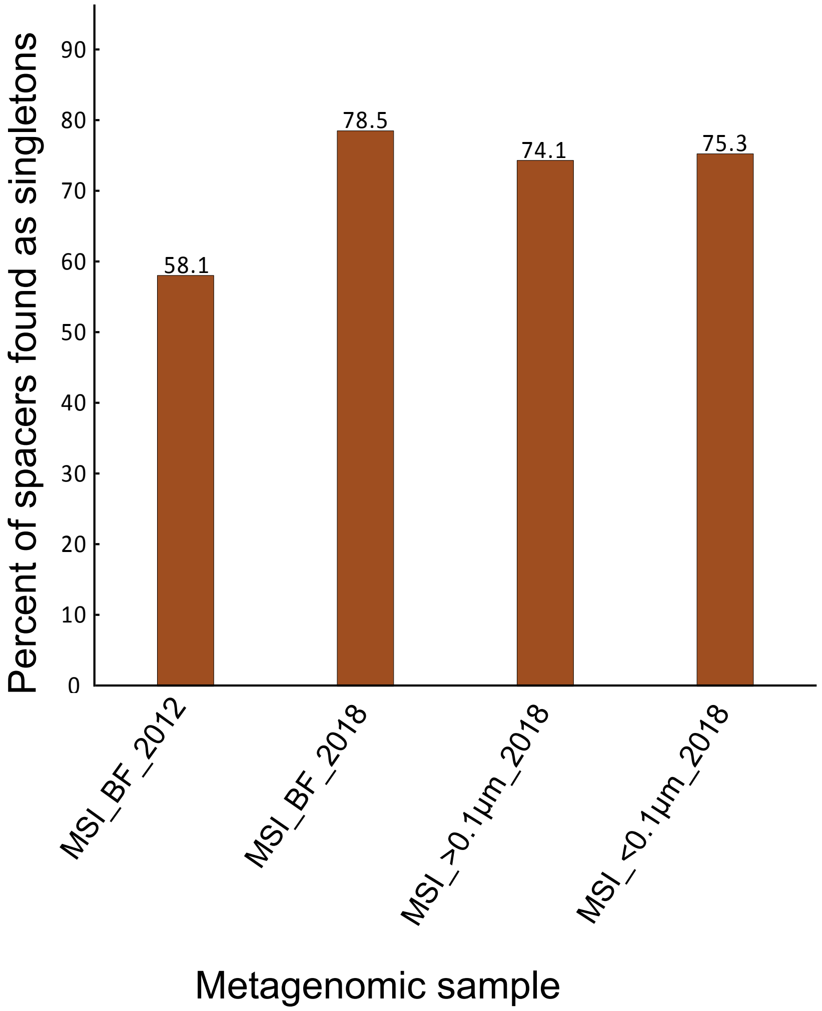

**Figure S11: GenomeFISH of a dense Altiarchaeota biofilm (BF) flock showing no detectable infection by Altivir_1_MSI.** Based on our enumeration, the majority of Altiarchaeota BF were infected by Altivir_1_MSI, however two out of seventeen BF flocks hybridized with the Altivir_1_MSI probe mix showed no infections at all. BF material was analyzed with A) DAPI (blue, cells), B) ATTO 488 (green, 16S rRNA signal) and C) Alexa 594 (red, viral genomes of Altivir_1_MSI) and merged (D).

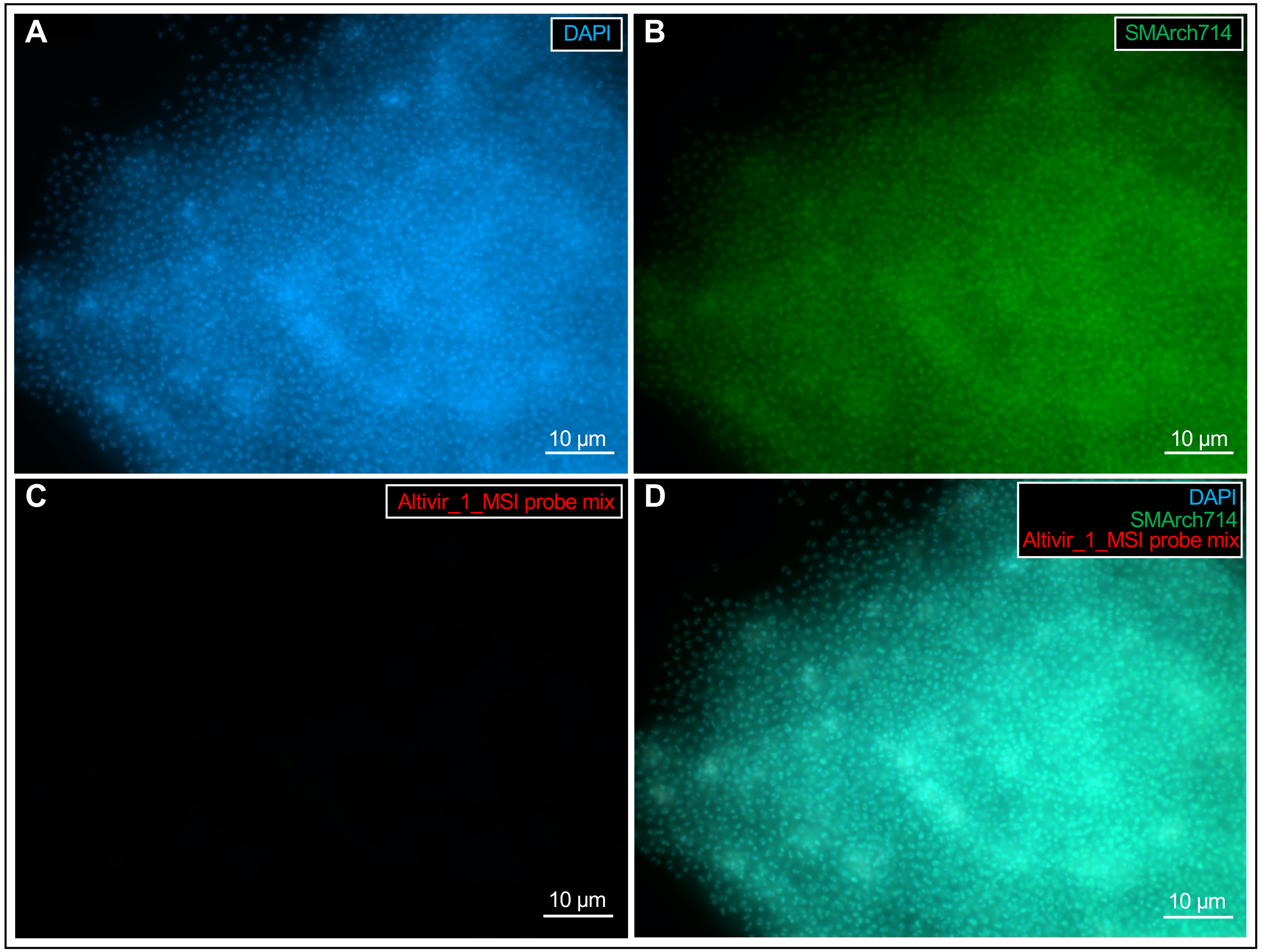

**Figure S12: EPI-fluorescence micrograph illustrating a heavily infected flock caused by Altivir_1_MSI.**
For genomeFISH, a chemically synthesized probe mix targeting Altivir_1_MSI was used. Biofilm material was analyzed with A) DAPI (blue, cells), B) ATTO 488 (green, 16S rRNA signal) and C) Alexa 594 (red, viral genomes of Altivir_1_MSI) and merged.

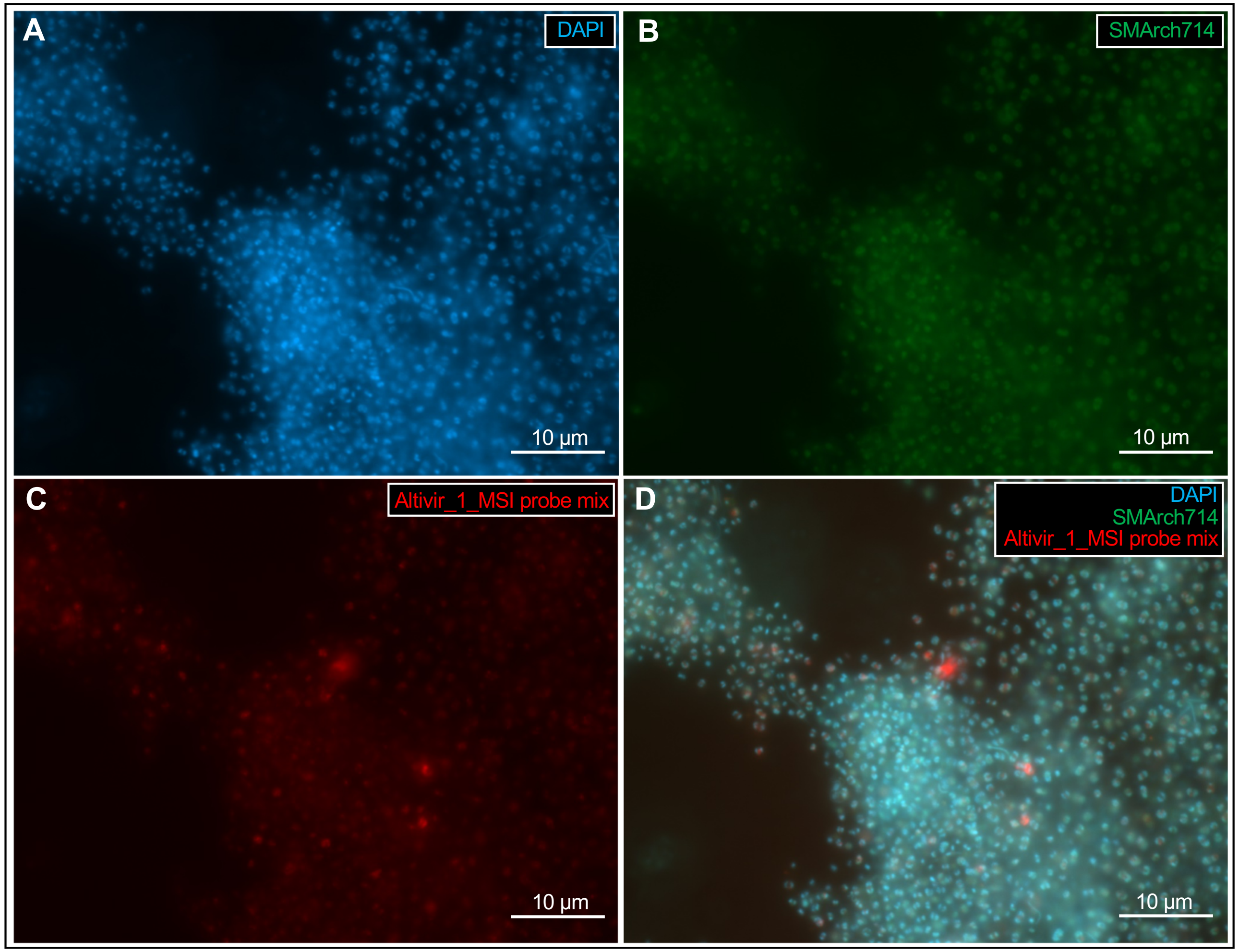

**Figure S13: EPI-fluorescence micrograph showing a heavily infected flock containing some rod-shaped bacteria potentially benefitting from lysing Altiarchaeota cells.** For genomeFISH, a chemically synthesized probe mix targeting Altivir_1_MSI was used. Biofilm material was analyzed with A) DAPI (blue, cells), B) ATTO 488 (green, 16S rRNA signal) and C) Alexa 594 (red, viral genomes of Altivir_1_MSI) and merged (D).

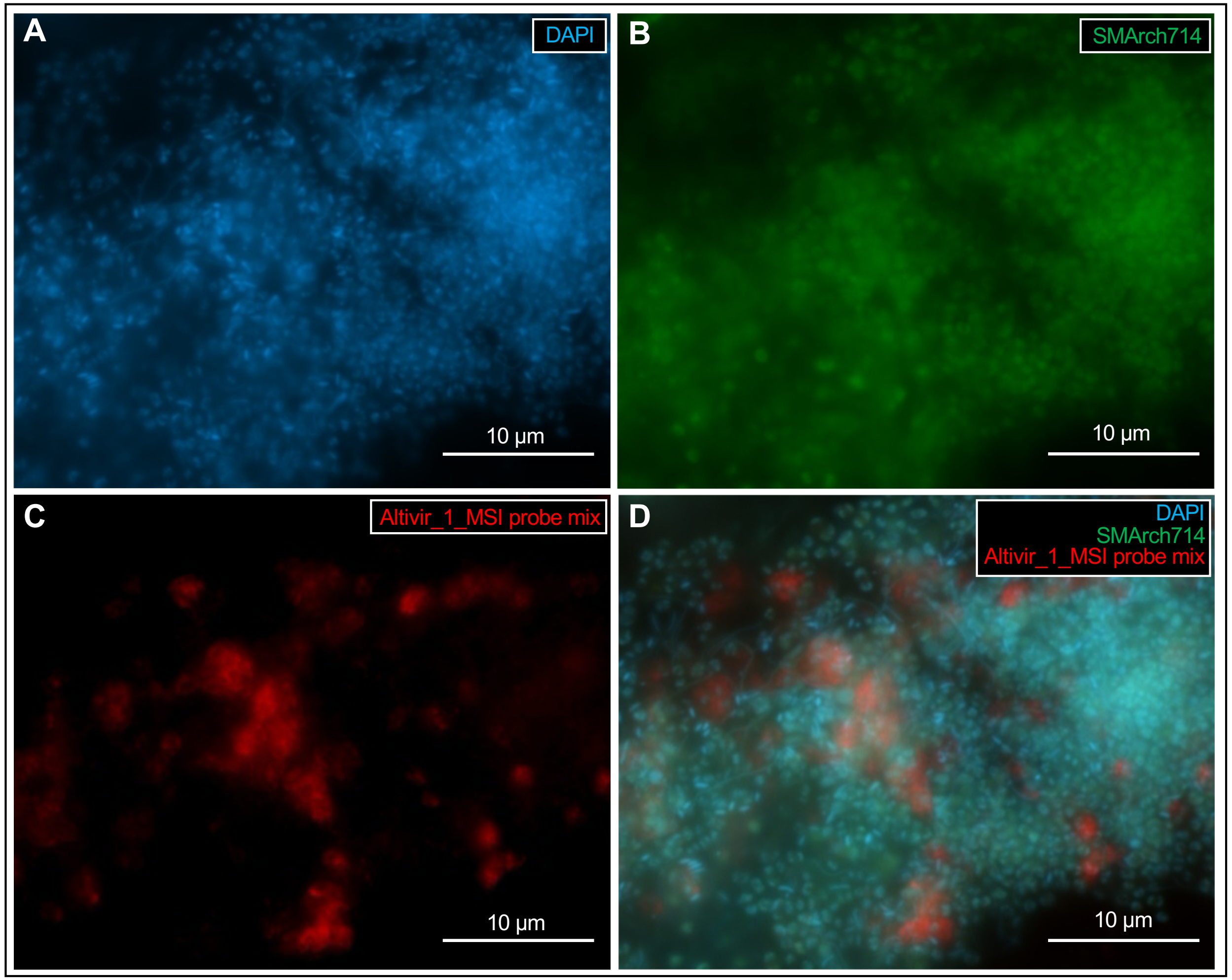
